# Efficient and Versatile Rapeseed Transformation for New Breeding Technologies

**DOI:** 10.1101/2024.11.06.622292

**Authors:** Kea Ille, Siegbert Melzer

**Author notes:** There are no data used in the manuscript that have to be stored in repositories.

## Abstract

Many gene functions are widely studied and understood in Arabidopsis; however, the lack of efficient transformation systems often limits the application and verification of this knowledge in crop plants. *Brassica napus* L., a member of the Brassicaceae family, is usually transformed by *Agrobacterium*-mediated hypocotyl transformation, but not all growth types are equally amenable to transformation. In particular, winter-type rapeseed, which requires vernalization to initiate flowering, is recalcitrant to *in vitro* regeneration and transformation. The analysis of gene functions in rapeseed is further complicated by the allotetraploid nature of its genome and the genome triplication within the *Brassica* genus, which has led to the presence of a large number of gene homologs for each Arabidopsis ortholog. We have established a transformation method that facilitates the regeneration of winter-type rapeseed by using the *WUSCHEL* gene from *Beta vulgaris*. This allowed us to efficiently transform winter-type and spring-type rapeseed in small-scale experiments. As proof of principle, we targeted *Bna.CLV3* and *Bna.SPL9/15* with CRISPR/Cas9 and showed that entire gene families are effectively edited using this transformation protocol. This allowed us to simultaneously study many redundantly acting homologous genes in rapeseed. We observed mutant phenotypes for *Bna.CLV3* and *Bna.SPL9/15* in primary transformants, indicating that biallelic knockouts were obtained for up to eight genes. This allowed an initial phenotypic characterization to be performed already a few months after starting the experiment.

## Introduction

Genetic variation is the driving force for establishing new traits in crop plants. New breeding techniques that employ gene editing are developing rapidly and offer many new methods to generate genetic variation. However, the transformation and regeneration of many crop plants is still a bottleneck for applying these techniques and for studying gene functions by molecular genetic approaches. This is even more challenging in polyploid crops that require efficient techniques to study highly redundant gene functions of many genes involved in a certain trait. Monocot crops, or crops such as sugar beet, are considered recalcitrant to *Agrobacterium*-mediated transformation, and recovering transgenic plants from these crops is difficult or impossible.

The improvement of crops, including the allotetraploid crop rapeseed (*Brassica napus* L.), with new breeding techniques is under constant development. As an important oilseed crop, rapeseed is cultivated worldwide and is adapted to distinct environments by different life history traits. Spring types can be sown and harvested within the same year, whereas semi-winter and winter types are biennial and require a mild or strict cold period (vernalization). *B. napus* is closely related to the model plant Arabidopsis; however, transformation and genetic engineering are not as straightforward as in Arabidopsis (Clough and Bent, 1988). Although some studies have performed *Agrobacterium*-mediated transformation of *B. napus* through floral dipping (Wang et al., 2003; Verma et al., 2008; Li et al., 2010; Ren et al., 2012), it is not widely and easily applicable. Therefore, most transgenic rapeseed plants have been generated through tissue-culture approaches. In tissue culture, the tissue type plays an important role in the success of the transformation and various tissues have been used, including haploid microspore-derived embryos (Boutilier et al., 2002), epicotyls, and higher stem segments (Cao Chu et al., 2020). However, most studies have used either cotyledons (Moloney et al., 1989; Bhalla and Singh, 2008) or hypocotyls (Radke et al., 1988; De Block et al., 1989; Cardoza and Stewart, 2003) for transgenic studies. In addition to culture conditions, medium composition and the concentration of phytohormones, *Agrobacterium* strains and densities are critical for transformation success. But even if rapeseed is generally amenable to transformation, protocols are often still genotype dependent, and not all relevant genotypes can be successfully transformed. The spring-type cultivar Westar is mostly used, whereas other studies also report the transformation of different spring types and some semi-winter-type rapeseed. Recent reports about transgenic winter-type rapeseed are lacking and although some early studies have demonstrated transformation of winter-type rapeseed (De Block et al., 1989; Damgaard et al., 1997; Chandler et al., 2005), it appears to be mostly recalcitrant to transformation. However, to be able to study winter-type-specific traits such as vernalization requirement, it is still important to generate mutants. Therefore, improving the transformation and regeneration of winter-type rapeseed in particular, is crucial.

Recently, the efficiency of regeneration and transformation of other recalcitrant crop species or genotypes could be improved by the expression of morphogenic genes. The transformation of wheat has been improved by using wheat *GROWTH-REGULATING FACTOR 4* (*GRF4*) and its cofactor *GRF-INTERACTING FACTOR 1* (*GIF1*) (Debernardi et al., 2020) and *WUSCHEL-RELATED HOMEOBOX 5* (*WOX5*) (Wang et al., 2022). Moreover, the transformation rates of recalcitrant sugar beet varieties, soybean, sunflower, and maize could be enhanced by overexpression of *GRF5* (Kong et al., 2020). *WUSCHEL* (*WUS*) belongs to the same homeobox gene family as *WOX5* and has been widely used to improve transformation and regeneration. Two of the first morphogenic genes used for transformation enhancement were *ZmWUS2* and *ZmBBM* (*BABYBOOM*), which enabled the transformation of previously non-transformable maize inbred lines and other crops including sorghum (Lowe et al., 2016); however, an excision system or specific promoters (Lowe et al., 2018) were necessary to avoid pleiotropic effects. Hoerster et al. (2020) used the cell non-autonomous nature of WUS proteins and developed an altruistic transformation strategy for maize in which the ZmWUS2 protein initiates somatic embryogenesis of neighboring cells transformed with a gene of interest without *ZmWUS2* integration.

However, the use of morphogenic genes for regeneration and transformation improvement in rapeseed is limited. It was shown that the overexpression of *BnBBM* in rapeseed caused the formation of somatic embryos on shoot tissues (Boutilier et al., 2002), and overexpression of *B. napus SHOOTMERISTEMLESS* (*STM*) increased the yield of microspore-derived embryos (Elhiti et al., 2010). Kong et al. (2020) also demonstrated that the overexpression of *AtGRF5* and *BnGRF5-LIKE* increased the amount of transgenic callus but did not contribute to the formation of transgenic shoots. To date, no transformation protocols for the use of morphogenic genes to improve rapeseed transformation are available, despite the lack of transgenic winter-type rapeseed.

Here, we present the use of the *WUS* ortholog from *Beta vulgaris* in *Agrobacterium*-mediated rapeseed hypocotyl transformation of *B. napus*. The overexpression of *BvWUS* facilitated the regeneration of transgenic winter-type rapeseed Express617 plants and enabled us to develop an easy, rapid, and small-scale applicable protocol. Furthermore, we observed CRISPR/Cas9 biallelic editing of multiple gene copies with redundant functions already in primary Express617 and Westar transformants, which allows the early analysis of gene functions.

## Results

### *BvWUS* facilitates shoot regeneration in winter-type rapeseed Express617

The overexpression of plant developmental regulators has facilitated and improved the regeneration of different recalcitrant plant species. Therefore, we initially aimed for a simple and efficient transformation protocol for sugar beet, known to be recalcitrant to *Agrobacterium*-mediated transformation. We tested whether the regeneration capacity and the transformation rate could be improved by expressing the Arabidopsis *WUS* gene and its ortholog from sugar beet. For this, we cloned the respective cDNAs into the binary vector p9o-LH-35s-ocs (DNA Cloning Service, Hamburg) under the control of the CaMV 35S promoter (35S::*AtWUS* and 35S::*BvWUS*) and used these constructs for *Agrobacterium*-mediated transformation of different tissues. However, none of our experiments gave rise to fully developed transgenic sugar beet plants.

In parallel, we were developing a regeneration and transformation strategy for the recalcitrant winter-type rapeseed Express617. We thus tested whether the expression of either *AtWUS* or *BvWUS* under the control of the CaMV 35S promoter would improve the regeneration and transformation efficiency in Express617 and give rise to transgenic plants. Accordingly, we carried out a small-scale hypocotyl transformation with the 35S::*AtWUS* and 35S::*BvWUS* constructs using 295 and 282 hypocotyl segments, respectively. After three weeks, hypocotyls transformed with the 35S::*AtWUS* construct showed massive embryogenic growth at their ends. Numerous somatic embryos formed from the callus and over time, the hypocotyls progressively produced more embryogenic structures and substantial swelling of the callus (Figure 1a). However, none of these embryos fully developed into shoots.

**Figure 1.**
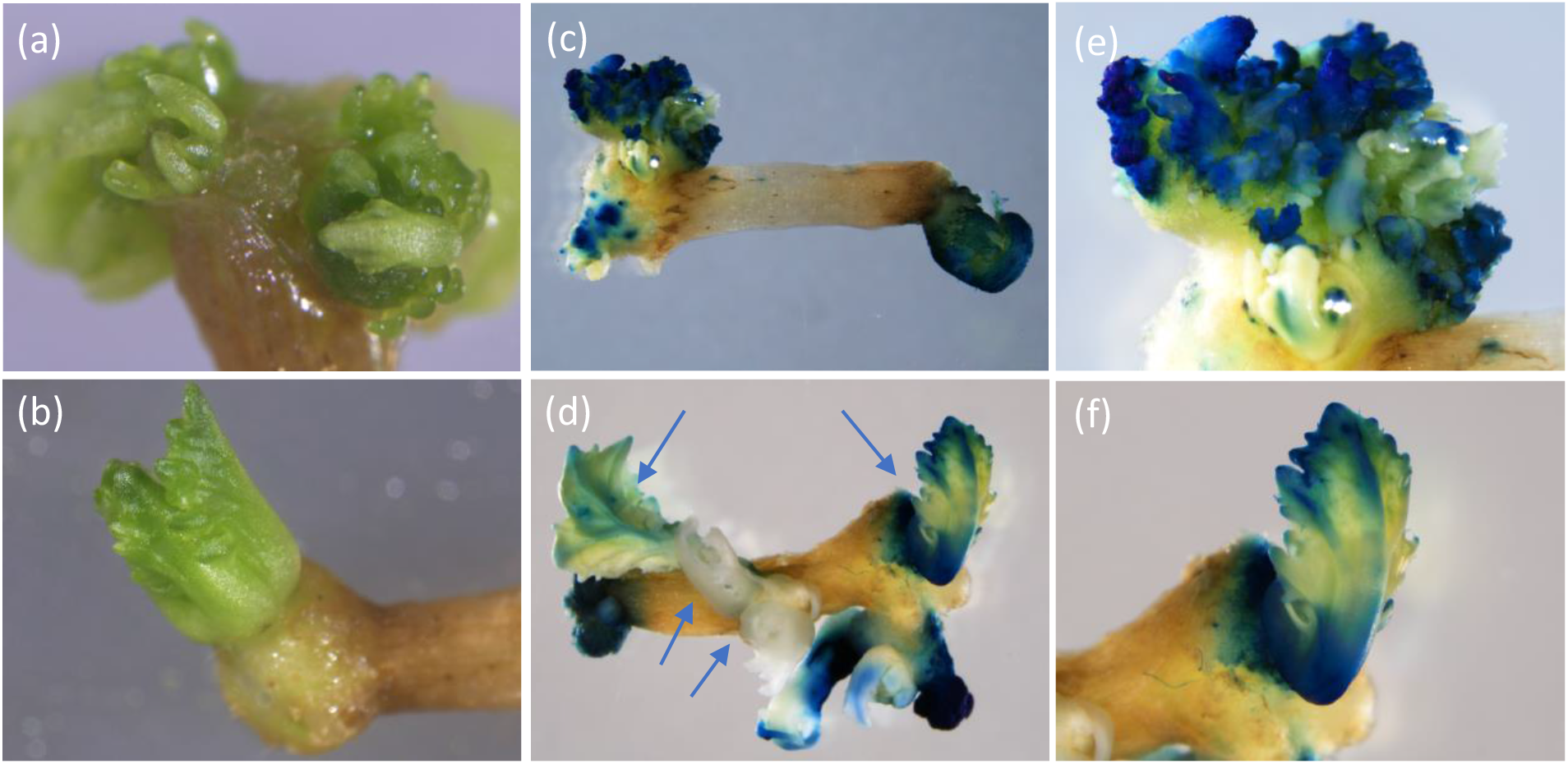
Express617 hypocotyl on selection medium after transformation with (a) 35S::*AtWUS*. Overproliferation of somatic embryos on hypocotyl ends without proper shoot formation; (b) 35S::*BvWUS*. Distinct single shoots grew from the callus at the hypocotyl end. (c) 35S::*AtWUS* and 35S::*GUS*. Blue-stained and white somatic embryos on hypocotyl ends after GUS staining. (d) 35S::*BvWUS* and 35S::*GUS*. Blue-stained shoots on hypocotyl after GUS staining. Destained green shoots appear yellowish whereas white shoots bleached out due to kanamycin selection. (e) Close-up of (c). (f) GUS-stained shoot from (d).

By contrast, hypocotyls transformed with the 35S::*BvWUS* construct developed many individual and distinct shoots directly from calli at the hypocotyl ends after six weeks (Figure 1b). In total, we regenerated 115 shoots, most of which subsequently bleached out under kanamycin selection. Nevertheless, 10 shoots on rooting medium developed roots and contained the *BvWUS* transgene. This demonstrates that expression of the distantly related *BvWUS* (Figure S1) in winter-type rapeseed hypocotyls can improve regeneration and transformation efficiency.

### Co-transformation with *BvWUS* can lead to transgenic plants that only contain the gene of interest

On the basis of the observation that most of the regenerated 35S::*BvWUS* shoots bleached out under kanamycin selection, we reasoned that due to the cell non-autonomous activity of WUS proteins (Daum et al., 2014), WUS might have moved from transgenic cells into neighboring cells lacking the transgene. Therefore, WUS proteins initiated shoot regeneration, and thus, many shoots regenerated that did not contain the transgene and the selection marker. On the basis of this assumption, we tested whether employing a co-transformation strategy with 35S::*AtWUS* and 35S::*BvWUS* together with another construct containing a gene of interest would facilitate the regeneration of plants not only harboring the *WUS* transgene and the gene of interest, but also of plants only containing the gene of interest. A high proportion of regenerated plants containing only the gene of interest would later eliminate the need to identify segregants without the *WUS* transgene.

Therefore, we mixed *Agrobacterium* containing the binary plasmid pBin19::pSH4 (35S::*GUS*) (Holtorf et al., 1995) and *Agrobacterium* with 35S::*AtWUS* or 35S::*BvWUS* in ratios of 9:1 and 1:1, respectively, and carried out co-transformation of Express617 hypocotyls. Similarly to in the transformation experiment with *WUS* only, hypocotyls co-transformed with 35S::*GUS* and 35S::*AtWUS* showed many somatic embryos (Figure 1c, e), and hypocotyls co-transformed with 35S::*GUS* and 35S::*BvWUS* showed *de novo* shoot formation (Figure 1d, f). To determine the integration of the 35S::*GUS* transgene, we performed a GUS assay on hypocotyls with regenerated tissues.

Hypocotyls co-transformed with 35S::*GUS* and 35S::*AtWUS* showed numerous GUS-stained embryos and some areas of developing callus and embryos without any GUS staining (Figure 1c, e). Hypocotyls co-transformed with 35S::*GUS* and 35S::*BvWUS* showed three types of shoots. Some shoots were white because they were already bleached out under kanamycin selection, indicating that no T-DNA was inserted. Some green shoots did not show any GUS staining and were identifiable as shoots with a yellowish appearance after destaining with ethanol, which suggests integration of the *BvWUS* transgene alone. Green shoots that turned blue in the GUS assay indicated the integration of the *GUS* transgene alone, or the co-integration of both transgenes (Figure 1d).

From the co-transformation experiment with 35S::*GUS* and 35S::*BvWUS*, we recovered 54 transgenic plants, of which 24 contained the GUS transgene and 19 were double transformants with both transgenes integrated. The other 5 plants were sole *GUS* transformants and 30 plants contained only the *WUS* transgene. The co-transformation experiment with 35S::*GUS* and 35S::*AtWUS* resulted in only one transgenic plant (Table 1).

**Table 1:**
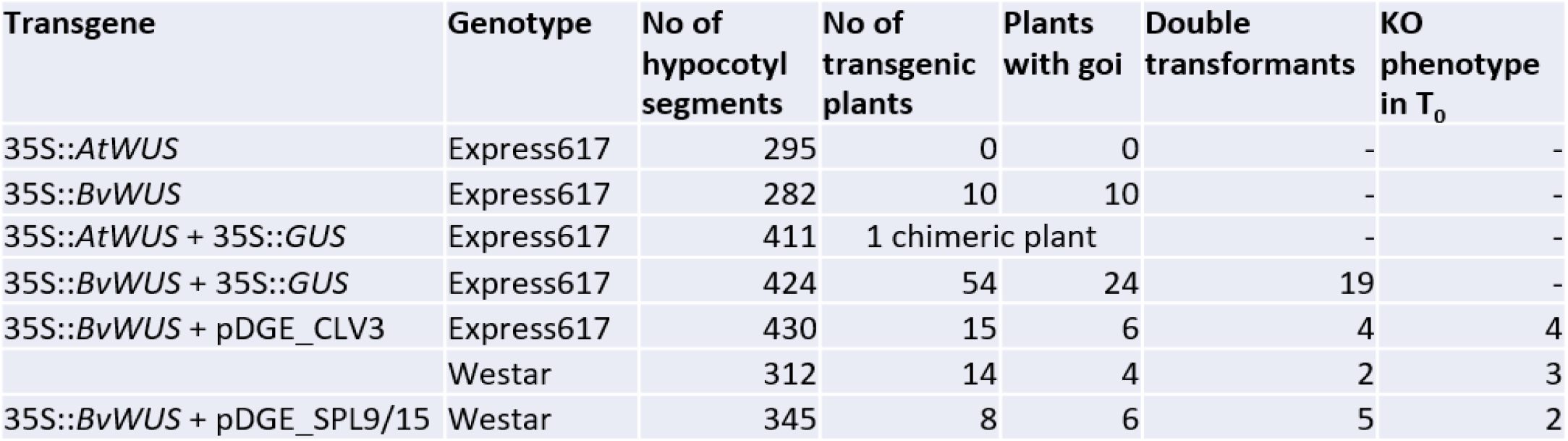
Overview of transformation experiments and obtained transgenic T_0_.

We observed that plants containing the *BvWUS* transgene showed a specific phenotype: In tissue culture and shortly afterwards, 35S::*BvWUS* plants often produced multiple shoots from the plant base (Figure 2a), which after vernalization (Figure 2b) all developed inflorescences. We further tested whether the phenotype was also transmitted to the T_1_ generation. However, none of the T_1_ progeny produced more than one shoot (Figure 2c), and no other mutant phenotypes.

**Figure 2.**
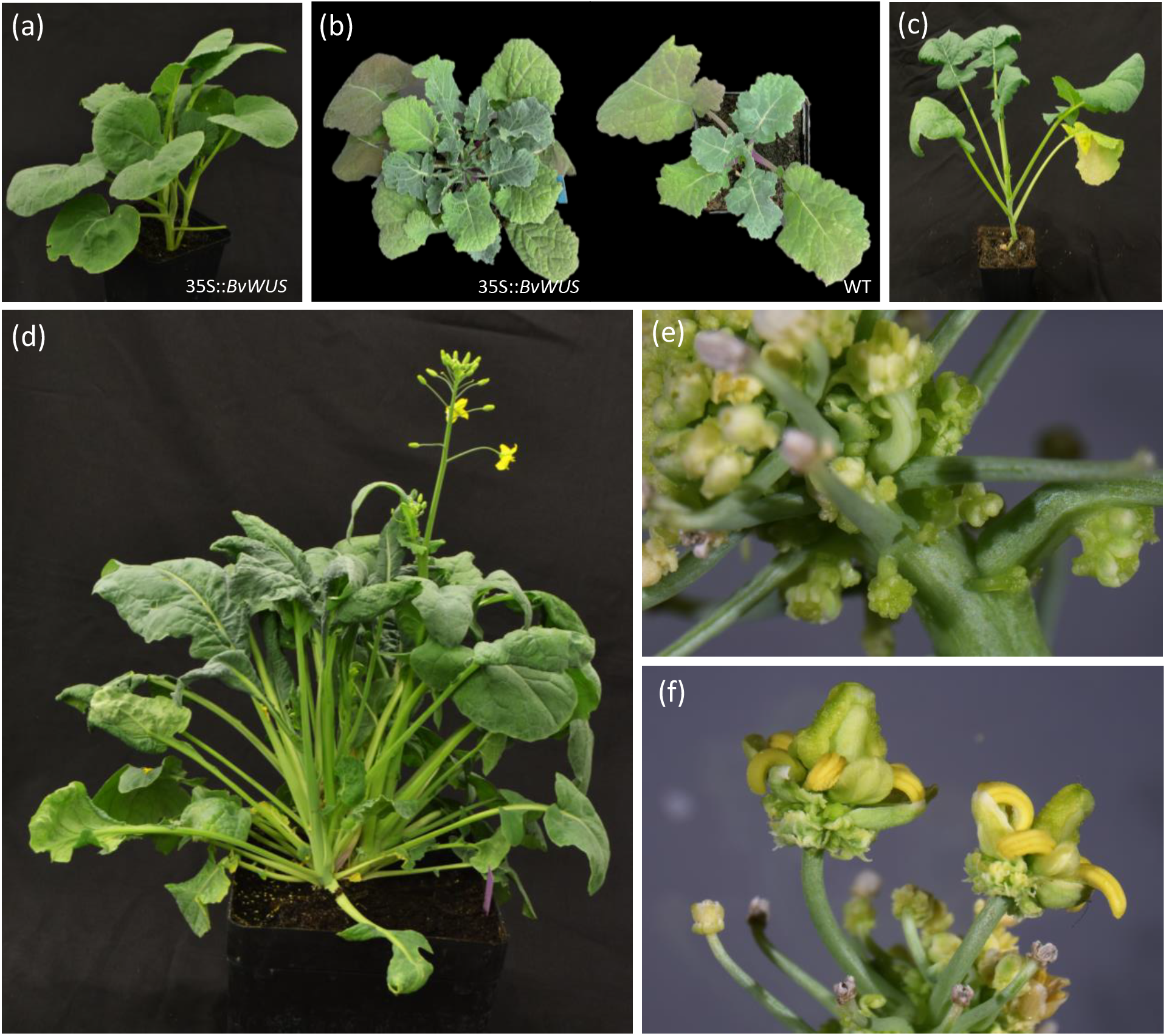
Express617 plants expressing *BvWUS* or *AtWUS*. (a) T_0_ 35S::*BvWUS* plant shortly after transplanting to soil has two shoots forming from the base of the plant. (b) T_0_ 35S::*BvWUS* plant and WT plant right after vernalization with multiple shoots and one shoot, respectively. (c) 35S::*BvWUS*T_1_ plant with one shoot. (d) Chimeric 35S::*AtWUS* plant at flowering stage. (e) Somatic embryos on the inflorescence stem on the chimeric part of the 35S::*AtWUS* plant. (f) Misshapen flowers of a chimeric 35S::*AtWUS* T_0_ plant.

Notably, the single *AtWUS* T_0_ plant also produced multiple shoots (Figure 2d), similarly to 35S::*BvWUS* plants. One of the shoots had crinkled leaves and along the inflorescence there was the massive development of embryogenic structures that were comparable to the embryogenic growth we observed on regenerating calli on hypocotyls (Figure 2e), whereas other shoots developed normally. This indicates that only that particular shoot contained the 35S::*AtWUS* transgene, whereas the remaining shoots had a WT phenotype, implying that the regenerated plant was chimeric. Flowers on the 35S::*AtWUS* shoot were misshapen with degenerated floral organs and consequently did not produce any seeds (Figure 2f).

These experiments showed that using a co-transformation strategy with 35S::*GUS* and 35S::*BvWUS* can lead to the recovery of mostly co-transformed plants that contain both transgenes, and plants that only contain the 35S::*BvWUS* transgene. However, a small number of plants with only the gene of interest can also be recovered, although these plants are underrepresented among the regenerated transgenic plants. Moreover, the expression of *BvWUS* did not alter the phenotype of the plants, indicating that mutant phenotypes could potentially be observed even in the presence of the transgene. By contrast, the expression of *AtWUS* only rarely facilitated the recovery of shoots and had detrimental effects in the recovered chimeric plant, such as the overproliferation of somatic embryos and infertility. Hence, expressing *AtWUS* was not suitable for improving the transformation and regeneration of rapeseed.

### Co-transformation with *BvWUS* and CRISPR vectors can lead to the efficient editing of multiple *BnaCLV3* alleles

Next, we aimed to establish a 35S::*BvWUS* and CRISPR/Cas9 co-transformation strategy for gene editing. As proof of principle, we chose to knock out both *Bna.CLAVATA3* (*CLV3*) homologs (*Bna.CLV3.A04* and *Bna.CLV3.C04*) in the winter-type rapeseed Express617 and spring-type rapeseed Westar. As previously demonstrated (Yang et al., 2018), *bna.clv3* mutants showed a clear phenotype with increased leaf number, an enlarged SAM, and multilocular siliques, which fulfilled our requirement for a phenotype that was easy to detect. We targeted both *Bna.CLV3* genes with four sgRNAs (Figure 3a) cloned in the plant transformation vector pDGE652 (zCas9i cloning kit, Stuttmann et al., 2021). Target sites TS1 and TS4 were conserved between *Bna.CLV3.A04* and *Bna.CLV3.C04*. TS2 (C04), and TS3 (A04) differed by one SNP 7 bp upstream of the PAM site (Table S1). This resulted in three different target sites per allele and thus a total of 12 target sites in all four *Bna.CLV3* alleles.

**Figure 3.**
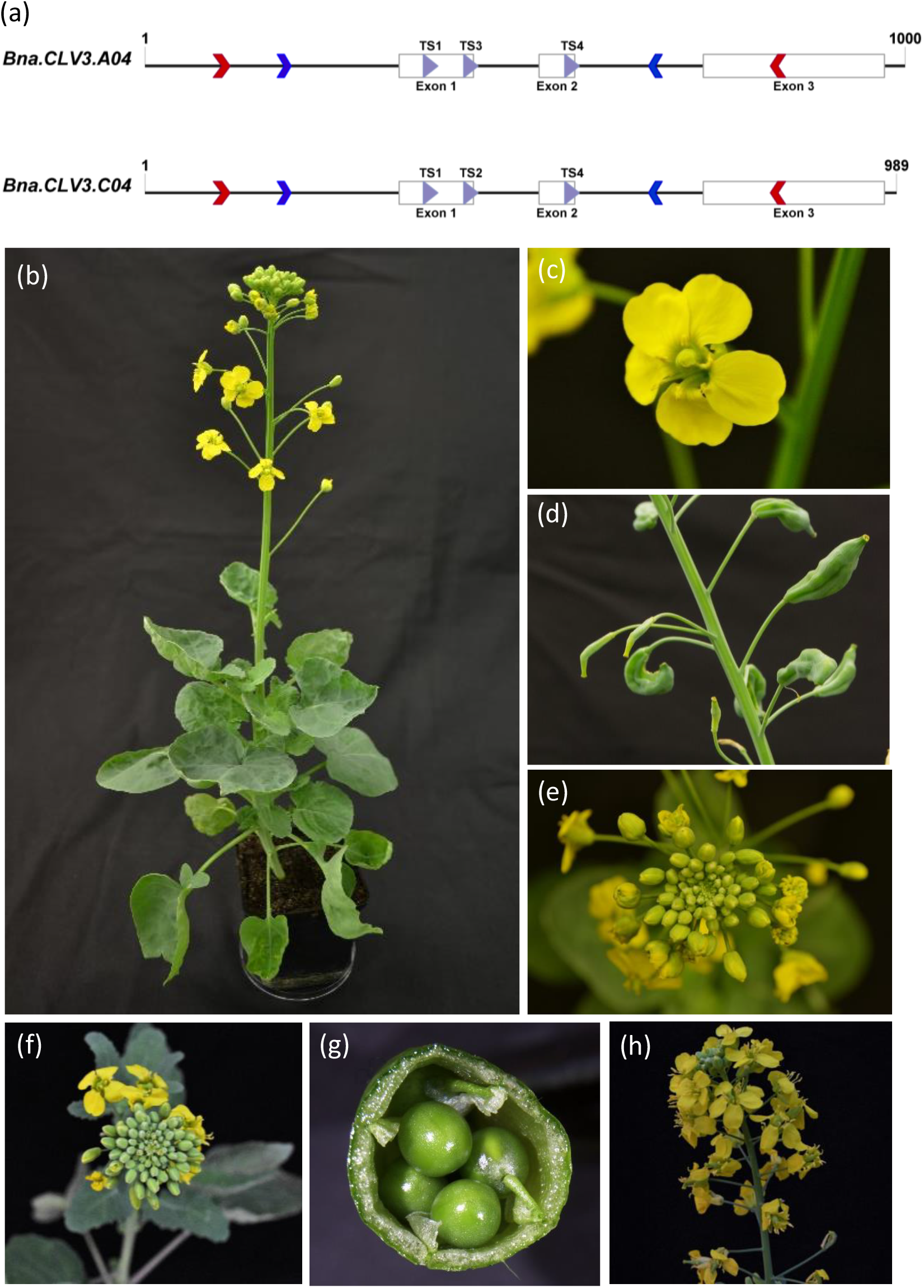
*Bna.CLV3* editing in Westar. (a) Gene structure of *Bna.CLV3.A04* and *Bna.CLV3.C04*. Blue arrows indicate the position of conserved primers with barcodes for amplicon sequencing. Red arrows show the position of homolog-specific primers and purple arrows show the position of target sites. (b) T_0_ Westar plant 7 with *bna.clv3* full knock-out phenotype. (c) Flower of W7 with five petals and enlarged gynoecium. (d) Multilocular silique of W7. (e) Expanded shoot apical meristem (SAM) in W7. (f)–(h) bna.clv3 full knockout phenotype in progeny of W7 (T_1_) with (f) expanded SAM, (g) multilocular silique, and (h) increased number of petals.

We carried out a small-scale 35S::*BvWUS* co-transformation with 430 and 312 hypocotyl segments of Express617 and Westar, respectively. The transformation of Express617 gave rise to 15 transgenic plants, of which 6 contained the CRISPR T-DNA whereas 4 were double transformants. Moreover, we obtained 14 transgenic Westar plants, of which 4 had the CRISPR T-DNA integrated, and 2 were double transformants (Table 1). Already in the T_0_ generation, we observed 4 and 3 plants with a *bna.clv3* knockout phenotype in Express617 and Westar, respectively. T_0_ *bna.clv3* plants showed an increased number of leaves and petals, as well as enlarged gynoecia and multilocular siliques (Figure 3b-e). The presence of a knockout phenotype in a high proportion of T_0_ plants indicates efficient editing in all four alleles of *Bna.CLV3* in both winter- and spring-type rapeseed.

### Analysis of *Bna.CLV3* gene-editing events

We analyzed the gene-editing events of T_0_ plants by amplicon sequencing (NGS Amplicon-EZ, Azenta). Amplicon sequencing requires PCR fragments of up to 500 bp spanning the target sites; however, designing gene-specific primers of homologous genes in such a small nucleotide range in rapeseed is challenging due to its allotetraploid nature and consequent sequence similarities. Thus, we designed conserved primers to amplify both *Bna.CLV3* copies simultaneously, including regions with SNPs to distinguish between the two copies (Figure 3a). To pool samples for sequencing, we attached barcodes to the primers and multiplexed up to five samples. For demultiplexing reads, we first joined paired-end reads using fastp (Chen, 2023), and employed the grep command in Unix to search for barcodes at the 5’ and 3’ ends of the reads. We analyzed demultiplexed reads with CRISPResso 2.0 (Clement et al., 2019) in CRISPRessoPooled mixed mode and further validated results by checking the output bam files in Integrative Genome Viewer (Robinson et al., 2011). However, we observed a high proportion of PCR recombination, which can occur if very similar genes from gene families or polyploid organisms such as rapeseed are amplified simultaneously (Meyerhans et al., 1990). If the amplification of a template stops prematurely, the incompletely transcribed strand can serve as a primer for the other homolog in our case, which leads to chimeric PCR products. We frequently observed that SNPs from both *Bna.CLV3* gene copies were included within one read (Figure 4a). These chimeric PCR products prevented us from determining to which homologs the mutations belonged. To circumvent these difficulties, we developed a strategy to minimize PCR recombination. Therefore, we designed homolog-specific primers around the target sites for an initial PCR and subsequently conducted a nested PCR with barcode primers (Figure 3a), which minimized PCR recombination.

**Figure 4.**
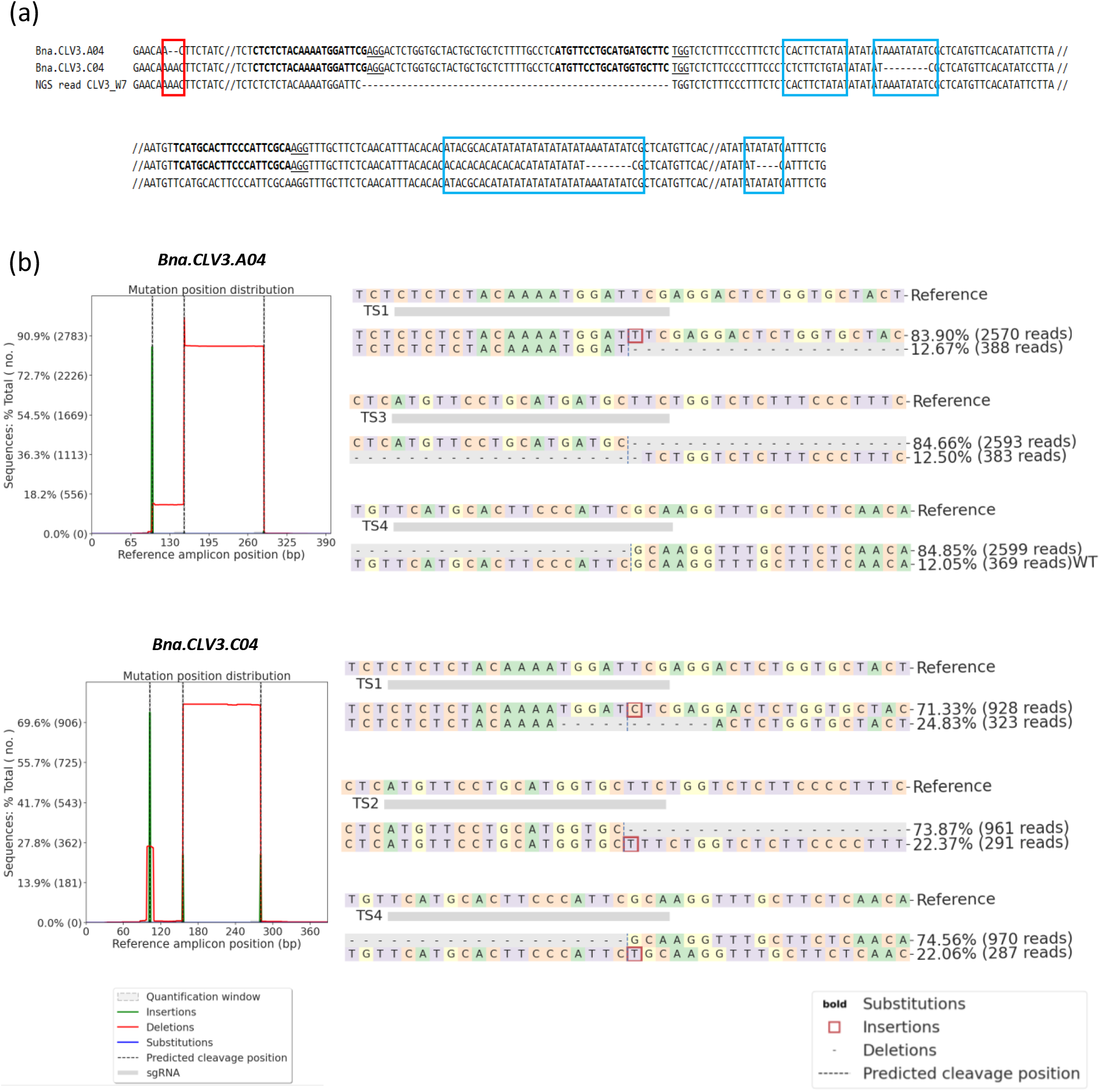
Analysis of gene editing in T_0_ plants by amplicon sequencing. (a) Example of PCR recombination when both gene copies are amplified together. The two upper rows display the reference sequence for *Bna.CLV3.A04* and *Bna.CLV3.C04*. Bold letters indicate the location of the target sites and underlined sequences represent the PAM site. The lowermost sequence is an example of a chimeric PCR product observed during amplicon sequencing. Red boxes indicate the affiliation of the SNP in the NGS read to homolog C04 whereas blue boxes indicate that the observed SNP matches the homolog A04. (b) CRISPResso2 results for *Bna.CLV3.A04* and *Bna.CLV3.C04* in T_0_ plant W7. Left: Frequency of insertions, deletions, and substitutions across the entire amplicon, considering only modifications that overlap with the quantification window. Right: Visualization of the distribution of identified alleles around the cleavage site for the TS1–TS4. Nucleotides are indicated by unique colors (A = green; C = red; G = yellow; T = purple). Substitutions are shown in bold font. Red rectangles highlight inserted sequences. Horizontal dashed lines indicate deleted sequences. The vertical dashed line indicates the predicted cleavage site.

In Westar plant 7 (*clv3_*W7), 11 out of 12 target sites were edited, resulting in a full knockout and a *bna.clv3* phenotype in T_0_. We identified biallelic editing in both genes, resulting in two different knockout alleles per gene. Collectively, four target sites (A04.1.TS1, C04.1.TS1, C04.2.TS2, and C04.2.TS4) showed an insertion of 1 bp, target site C04.1.TS1 showed a deletion of 11 bp and A04.2.TS4 maintained the WT sequence. We further observed large deletions of 133 bp between A04.1.TS3 and A04.1.TS4 and of 125 bp between C04.1.TS2 and C04.1.TS4, and another deletion of 54 bp between target sites A04.1.TS1 and A04.1.TS3 (Figure 4 b).

Out of 16 analyzed alleles of 4 transgenic Westar plants, 14 were knock-out alleles, and among 6 transgenic Express617 plants, 21 out of 24 analyzed alleles led to a knockout genotype. As indicated by the *bna.clv3* phenotype of seven T_0_ plants, all alleles and most target sites were edited in these plants. In three transgenic *bna.clv3* Westar and four Express617 plants, sequencing results confirmed the knockout phenotype and showed gene editing in all four alleles leading to a full knockout (Figure 5, Table S2). Overall, the number of target sites and edited alleles, and the frequent occurrence of large deletions between target sites indicated highly efficient gene editing using this *BvWUS* co-transformation strategy.

**Figure 5.**
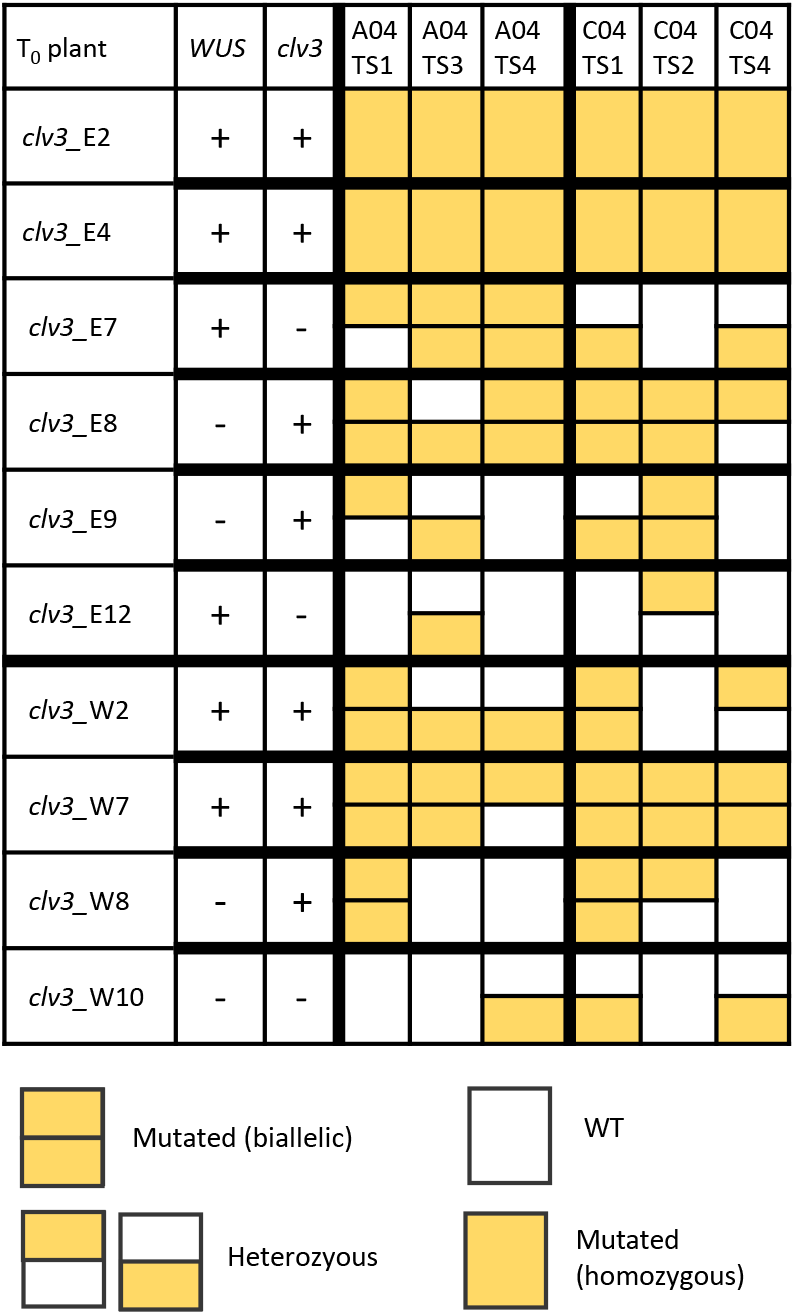
Genotyping of primary Westar and Express617 transformants from co-transformation with 35S::*BvWUS* and pDGE652_CLV3. Integration of *BvWUS* T-DNA was determined by PCR. The *bna.clv3* full knockout phenotype was recorded. Gene editing at each target site was analyzed by amplicon sequencing.

To obtain T-DNA-free T_1_ Westar plants and to check for the inheritance of the mutations and phenotype, we self-pollinated T_0_ Westar plants and sowed T_1_ seeds. Plants were screened for the *BvWUS* and CRISPR T-DNA by PCR. Among 48 T_1_ progeny of *clv3*_W2, we identified no plants that lacked either T-DNA, suggesting the presence of multiple integrations. Because W2 was edited biallelically in both genes, all progeny also showed the *bna.clv3* phenotype (Figure 3f-h). This was also the case for the progeny of *clv3*_W7 and *clv3*_W8, in which we determined the integration of one and no *BvWUS* transgene, respectively, and more than three CRISPR T-DNAs for *clv3*_W7 and 2 CRISPR T-DNAs for *clv3*_W8. The segregation pattern of the progeny of *clv3*_W10 indicated the integration of only one CRISPR T-DNA. Because the mutations in T_0_ were only heterozygous in A04.2.TS4, C04.2.TS1 and C04.2.TS4, the *bna.clv3* phenotype was absent and the progeny segregated for the *bna.clv3* knockout phenotype (Tables S3-6).

For Express617 we selected T_1_ seeds on the basis of red fluorescence of the FAST marker encoded within the T-DNA. Therefore, we germinated seeds on wet filter paper and screened seedlings for red fluorescence (Figure S2). T_1_ seeds without red fluorescence and thus without the CRISPR T-DNA were grown to check for the *bna.clv3* phenotype. All progeny of *clv3*_E2, *clv3*_E4, *clv3*_E8, and *clv3*_E9 showed the *bna.clv3* knock-out phenotype, whereas the progeny of *clv3*_E7 and *clv3*_E12 segregated for the phenotype.

In summary, all mutations and the *bna.clv3* knock-out phenotype were transmitted to the T_1_ generation and T-DNA-free plants could be identified.

### Eight *Bna.SPL9/15* genes can be efficiently edited simultaneously in Westar

Because we obtained high proportions of *bna.clv3* knockout phenotypes in T_0_ plants, and observed that most target sites were edited, we aimed to knock out a greater number of genes simultaneously. Therefore, we chose to target *Bna.SPL9* and *Bna.SPL15* in Westar, which both have four homologs in rapeseed, with six different sgRNAs. We designed target sites in the eight genes *Bna.SPL9.A04, Bna.SPL9.A05, Bna.SPL9.C04a, Bna.SPL9.C04b, Bna.SPL15.A04, Bna.SPL15.A07, Bna.SPL15.C04*, and *Bna.SPL15.C06*, targeting each gene once or twice (Figure 6a). TS1 is conserved between the genes *Bna.SPL15.A07* and *Bna.SPL15.C06*, and TS2 between the genes *Bna.SPL9.A05* and *Bna.SPL9.C04b*. TS3 from *Bna.SPL9.A04* has one mismatch on *Bna.SPL9.C04a* at position 18 upstream of the PAM site. TS4 is conserved between *Bna.SPL15.A07* and *Bna.SPL15.C06* and additionally has two mismatches at positions 19 and 20 upstream of the PAM site of *Bna.SPL15.A04* and one mismatch at position 20 of *Bna.SPL15.C04*. TS5 from *Bna.SPL9.C04b* has one mismatch at position 12 on *Bna.SPL9.A05*, and TS6 is conserved between *Bna.SPL9.A04* and *Bna.SPL9.C04a* (Table S1). In total, 14 sites in all 8 genes or 28 sites in 16 alleles were targeted.

**Figure 6.**
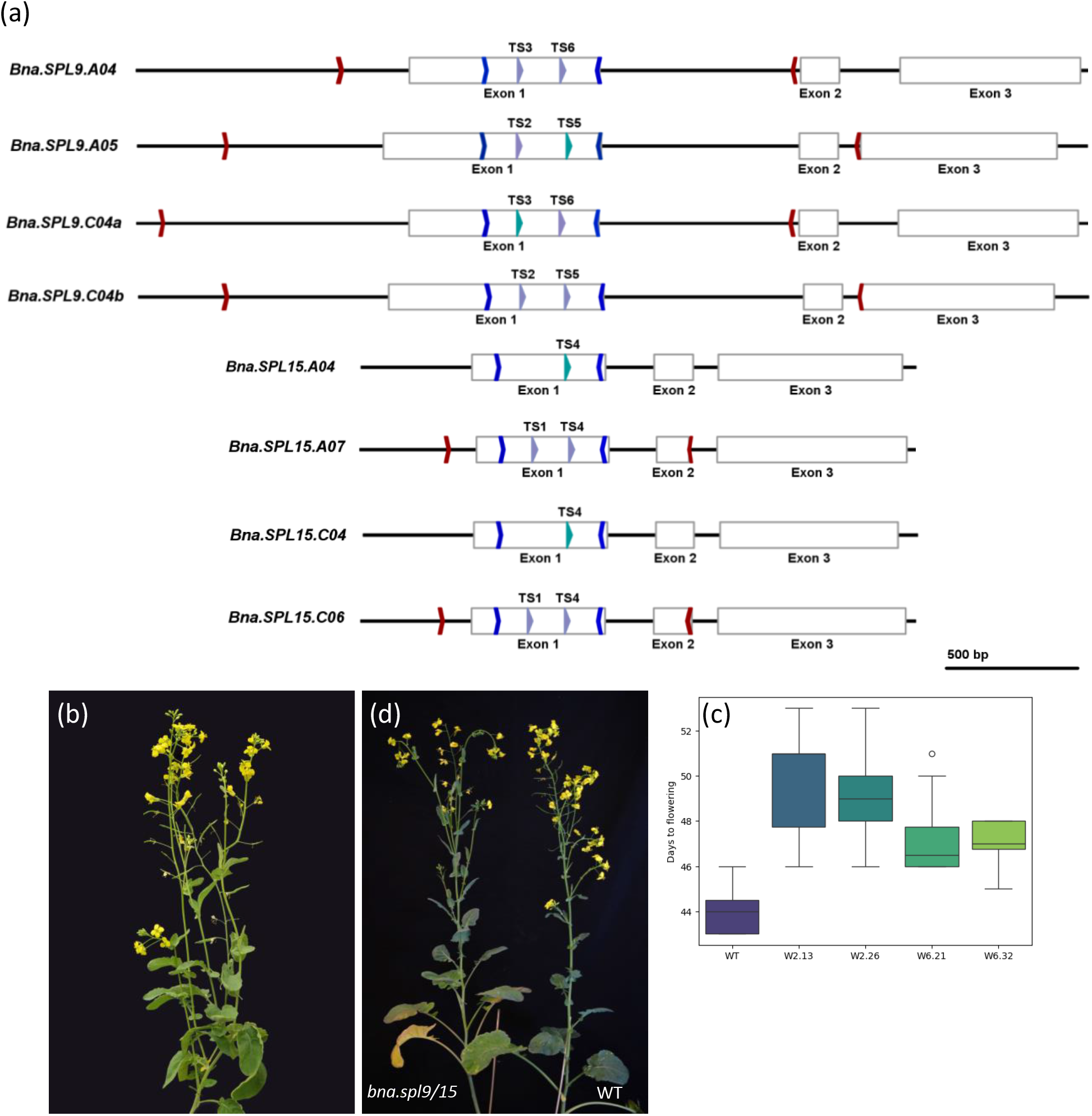
*BnaSPL9/BnaSPL15* editing in Westar. (a) Gene structure of *Bna.SPL9* genes and *Bna.SPL15* genes. Blue arrows indicate the position of conserved primers with barcodes for amplicon sequencing. Red arrows show the position of homolog-specific primers and purple arrows show the position of target sites. Turquoise arrows show the position of target sites with mismatches. (b) T_0_ *bna.spl9/15*_W2 with reduced apical dominance. (c) T_2_ *bna.spl9/15_*plant W6.32.6 with reduced apical dominance flowered later than Westar WT. (d) T_2_ families flowered later than Westar WT.

After the co-transformation of 345 Westar hypocotyl segments, we obtained eight transgenic plants of which six contained the CRISPR T-DNA. Furthermore, five of these plants also contained the 35S::*BvWUS* transgene and were thus co-transformants. Only one plant contained solely the CRISPR T-DNA (Table 1). *Bna.SPL9/15* T_0_ plants *spl9/15*_W2 and *spl9/15*_W8 showed a reduction in apical dominance (Figure 6b) as described for the *spl9 spl15* Arabidopsis mutant (Schwarz et al., 2008) suggesting a full knockout of all eight genes.

We sequenced *SPL9.C04a* and *b* in plants *spl9/15*_W1, *spl9/15*_W2, *spl9/15*_W4, and *spl9/15*_W6 and found large deletions between target sites TS2 and TS5 in *Bna.SPL9.C04b* in all four plants (Figure S3). We further observed editing of TS3 which had one mismatch on *Bna.SPL9.C04a* in all four analyzed plants. Notably, both alleles of *Bna.SPL9.C04a* in plant *spl9/15*_W2 were only edited at the target site with mismatches but not at the conserved target site TS6 (Figure S4). We analyzed one plant with a loss of apical dominance phenotype (*spl9/15*_W2) and one WT phenotype (*spl9/15_*W6) in more detail and sequenced all targeted genes of these plants. In *spl9/15*_W2, all genes were edited in both alleles including *Bna.SPL15.A04* and *Bna.SPL15.C04*, which only had targets with mismatches leading to a full knockout of all genes. Out of 28 target sites in 16 alleles, 21 were edited, whereas 7 remained WT. By contrast, in T_0_ plant *spl9/15*_W6 we observed that only TS1 on one allele of *Bna.SPL15.A07* and TS4 on one allele of *Bna.SPL15.A04* were not edited. We found mutations in 26 out of 28 target sites; however, due to the lack of editing in one allele of *Bna.SPL15.A07*, this gene was not knocked out. In addition, TS5 in *Bna.SPL9.A05* with a mismatch at position 12 was edited in both alleles (Table S7).

We sowed T_1_ seeds of both T_0_ plants *spl9/15*_W2 and *spl9/15*_W6, identified plants that lacked both T-DNAs, and sequenced selected plants (Table S8). We collected seeds and subsequently sowed the T_2_ generation from two T_1_ plants per T_0_ plant (*spl9/15*_W2.13, *spl9/15*_W2.26, *spl9/15*_W6.21, and *spl9/15*_W6.32), in which all eight genes were knocked out (Table S8). We grew 10 T_2_ plants per line and the Westar WT control at 20°C and 16 h light in a growth chamber. We recorded the opening of the first flower and the plant habitus. The flowering time of mutant lines ranged between 45 and 51 days after sowing, meaning that all mutant lines flowered later than the WT (Figure 6d). All lines showed a loss of apical dominance (Figure 6c).

Collectively, our experiments showed that using the distantly related *WUS* gene from sugar beet does not lead to overproliferation of embryonic tissues in transgenic plants. Instead, it leads to the formation of organized shoots that are indistinguishable those of WT plants. Moreover, a co-transformation strategy with *BvWUS* demonstrated that many genes and target sites can be effectively edited simultaneously; in this case, up to 8 genes and 26 target sites in 16 alleles.

## Discussion

The genetic analysis of growth-type-specific traits has been difficult in all growth types of allotetraploid rapeseed because a reliable protocol for winter-type rapeseed transformation was lacking. Most studies that produced transgenic winter rapeseed were performed around 30 years ago and were not reproduced in more recent studies, despite seemingly straightforward transformation protocols (De Block et al., 1989; Damgaard et al., 1997).

However, elaborate transformation procedures via e.g. hairy-root transformation with *Agrobacterium rhizogenes* (Grison et al., 1996) or multiple tissue-culture steps (Stefanov et al., 1994) were previously used to obtain transgenic winter- and spring-type rapeseed. The use of tissue materials such as mesophyll protoplasts (Wang et al., 2005) or haploid microspores (Cegielska-Taras et al., 2008) might have hindered the reproducibility of these studies.

Due to the lack of a winter-type rapeseed transformation protocol, *Bna.SVP* was edited and analyzed in semi-winter rapeseed (Ahmar et al., 2022), which does not have a strict vernalization requirement; however, studying *SVP* in winter-type rapeseed would be more informative regarding the vernalization response.

We have developed a co-transformation strategy for recalcitrant winter-type rapeseed with the morphogenic gene *BvWUS*. Hypocotyl transformation allowed the easy, rapid, and small-scale generation of CRISPR/Cas9 winter- and spring-type rapeseed mutants. Using this protocol, the function of multiple gene homologs and whole gene families specific to winter- and spring-type rapeseed can now be analyzed. Using 35S::*BvWUS* has enabled the transformation and regeneration of winter-type rapeseed Express617 and improved the transformation efficiency of spring-type Westar. Our co-transformation strategy is based on simple *Agrobacterium*-mediated hypocotyl transformation (De Block et al., 1989). Hypocotyls are easily accessible and simple to handle in comparison with other tissues. The maintenance of the tissue cultures is undemanding due to the minimal necessary steps because the medium is changed and shoots are excised every three weeks. Moreover, the amount of starting material needed is low and in our experiments, up to 430 hypocotyl segments (equivalent to approximately 90 seedlings) was sufficient to generate several transgenic Express617 and Westar plants and CRISPR full knockouts in the T_0_ generation. LaManna et al. (2022) reported using 520 seedlings for *Agrobacterium*-mediated hypocotyl transformation of spring-type Westar and recovered only four plants that carried the T-DNA. Furthermore, to obtain T_1_ seeds, the whole procedure took about 12 months. With our co-transformation protocol, T_1_ Express617 and Westar transgenic seeds can be obtained with minimal labor within 8 and 6 months, respectively (Figure S5).

The overexpression of genes such as *GRF*, the chimera *GRF*-*GIF*, and *WOX5* improved the transformation of recalcitrant sugar beet (Kong et al., 2020) and wheat (Debernardi et al., 2020; Wang et al., 2022), respectively. Using these genes was advantageous because they did not lead to any abnormal phenotypes; however, the use of *BnaGRF5* in rapeseed increased the amount of transgenic callus but not the number of transgenic plants (Kong et al., 2020).

Another commonly used morphogenic gene is *WUS*, but simply overexpressing *WUS* can lead to severe pleiotropic effects. Overexpression of the endogenous *WUS* gene in Arabidopsis under the control of the 35S promoter led to abnormal phenotypes in seedlings and impaired development (Zuo et al., 2002). A 17-β-estradiol-inducible system helped to regulate *WUS* expression by restricting it to a certain time period, which enabled the formation of somatic embryos only during a short time window and led to the normal development of regenerated plants. The overexpression of *WUS* from different species apart from Arabidopsis has shown similar effects. Overexpression of the endogenous *BrrWUSa* in turnip also resulted in sterile and abnormal plants, which was also overcome by an estradiol-inducible system (Liu et al., 2022). The same system also helped to regenerate transgenic white spruce (Klimaszewska et al., 2010). Another strategy to limit *WUS* expression is the excision of the *WUS* cassette. Combining *ZmBBM* and *ZmWUS2* without their removal before regeneration also led to aberrant phenotypes and sterile plants in maize (Lowe et al., 2016). Furthermore, the ectopic expression of *AtWUS* in species such as cotton also caused abnormal somatic embryos (Zheng et al., 2014).

In our case, the expression of *AtWUS* and *BvWUS* in winter-type rapeseed showed remarkable differences. As previously described for other plant species, the ectopic expression of *AtWUS* in rapeseed also led to an aberrant phenotype and sterility. Whole plants could only be recovered when they were chimeric, suggesting that the constitutive expression of *AtWUS* hindered the development of fertile plants from somatic embryos. However, instead of using elaborate induction or excision systems to limit *AtWUS* expression in rapeseed hypocotyls, expressing the distantly related *BvWUS* (Figure S1) was sufficient to directly induce shoot formation instead of somatic embryos, and the regeneration of transgenic plants. Although regenerated transgenic T_0_ plants occasionally produced multiple shoots from the plant base, subsequent generations never showed this phenotype and no other aberrant phenotypes were observed.

In our previous attempts to transform Express617 without 35S::*BvWUS*, we were hardly able to regenerate any shoots from transformed hypocotyls. Using the 35S::*BvWUS* construct, we initially observed that many shoots formed on the callus, but most bleached out under kanamycin selection. We reasoned that the non-transgenic shoots formed due to the movement of cell non-autonomous BvWUS proteins from cells carrying the *BvWUS* T-DNA, to non-transgenic cells. Similarly, the movement of ZmWUS2 has been exploited for altruistic transformation of maize, and the transient expression of *ZmWUS2* and the simultaneous transformation of another T-DNA enabled the regeneration of transgenic plants that lacked the *ZmWUS2* construct (Hoerster et al., 2020).

On this basis, we developed a co-transformation strategy in which we mixed two *Agrobacterium* strains: One containing the 35S::*BvWUS* expression cassette and the other containing a construct of interest e.g. GUS or CRISPR. Throughout all experiments, only a small proportion of regenerated transgenic plants contained solely the construct of interest and not the 35S::*BvWUS* cassette (between 17% and 50%). In *B. napus*, ZS6 co-transformation efficiencies of up to 81% were obtained when two *Agrobacterium* strains, each containing a different selection marker, were mixed in a ratio of 1:1 (Liu et al., 2020). In our case, most transgenic plants contained the 35S::*BvWUS* cassette with co-transformation efficiencies between 14% and 62.5%, indicating that the regeneration of shoots is more efficient when the 35S::*BvWUS* cassette is integrated. In the altruistic transformation of maize, the co-transformation efficiency was low due to the strong expression of genes encoding WUS2 and CRC transcription factors in plants containing the altruistic T-DNA and the subsequent inhibition of growth of these plants (Hoerster et al., 2020). We reason that the low similarity between the BvWUS and the endogenous BnaWUS proteins (Figure S1) enables the regeneration of phenotypically normal rapeseed plants, and prevents the overproliferation of somatic embryos and aberrant phenotypes that are observed when using the more similar AtWUS. Moreover, we observed relatively low expression from the vector p9o-LH-35s-ocs (data not shown), which might be caused by the combination of a strong 35S promoter with a weak OCS (OCTOPINE SYNTHASE) terminator (De Felippes and Waterhouse, 2023). It remains to be tested whether stronger expression of *BvWUS* is beneficial for regeneration or leads to similar overproliferation.

Due to its allotetraploidy, the *B. napus* genome contains multiple copies per gene. It is therefore necessary to target and knock out several genes simultaneously to study gene functions. Several studies showed that this is not always trivial. In semi-winter-types J9712 and J9707, two gene copies of *BnaIDA* and *BnaTT8*, respectively, were targeted by CRISPR/Cas9 (Zhai et al., 2020; Wu et al., 2022). In J9712, only 7 out of the 30 transgenic plants exhibited editing events and none of the plants was knocked out for both targeted *BnaIDA* genes in the T_0_ generation. Zhai et al. (2020) observed five plants with seeds that had a yellow seed coat and thus a full knockout of both *BnaTT8* genes in T_0_; however, out of 333 transgenic plants, only 48 plants were edited, and only 5 for all alleles of these 2 genes. The same semi-winter line was previously used for editing two *CLV3* genes (Yang et al., 2018). 494 T_0_ plants carrying the T-DNA were recovered and only 51 of these plants were edited and 8 had a knockout phenotype.

In our experiments targeting *CLV3* in Westar and Express617, we recovered four and six plants that carried the CRISPR T-DNA, respectively. All plants with the CRISPR T-DNA were edited, and many plants were full knockouts displaying the described *clv3* phenotype (4/6 Express617 and 3/4 Westar plants). We were further able to increase the number of genes that were edited simultaneously, as shown in our CRISPR experiment targeting *BnaSPL9* and *BnaSPL15* in Westar. Again, we identified gene editing in all six plants carrying the CRISPR T-DNA. In two plants, all eight targeted *BnaSPL9* and *BnaSPL15* genes were edited in both alleles, which resulted in a full knockout in T_0_. We often observed large deletions between two or three target sites in both CRISPR experiments, and no obvious differences between editing of different target sites. Moreover, our selected targets with mismatches were efficiently edited. Collectively, this indicates an exceptionally high efficiency of gene editing using the *BvWUS* co-transformation strategy.

Che et al. (2022) also observed that CRISPR/Cas9 gene editing was more efficient when they used *ZmWUS2* in the altruistic transformation of sorghum. They hypothesized that WUS activity might alter chromatin accessibility, and thus might enable more efficient gene editing in open chromatin regions, which might also be a reasonable explanation for our high editing efficiencies when using *BvWUS*.

Another reason or additional factor for efficient editing might be the choice of the CRISPR vector. The pDGE vectors (Stuttmann et al., 2021) were specifically designed for multiplex editing of genes. Stuttmann et al. (2021) regenerated transgenic *Nicotiana benthamiana* plants from *Agrobacterium*-mediated transformation that were mutated in all eight targeted genes, mostly biallelically, with an overrepresentation of homozygous mutations. This is also consistent with our observation in rapeseed where an unexpectedly high number of T_0_ plants displayed full knockout phenotypes in the presented experiments. However, we also observed similar editing efficiencies using other CRISPR vectors (pCAS9-TPC (Fauser et al., 2014) and pAGM65881 (Grützner et al., 2021), unpublished data).

For ease of use and to circumvent the removal of two transgenes for CRISPR/Cas9, we integrated the 35S::*BvWUS* cassette into a CRISPR vector. Therefore, we modified the pCAS9-TPC vector extensively by changing the promoter of the *Cas9* gene, and integrating a kanamycin selection marker and a Golden Gate cassette. Preliminary experiments that targeted *CLV3* in Express617 suggested that transgenic, gene-edited plants with a full knock-out *clv3* phenotype can be obtained using this vector (data not shown).

In summary, we have developed an efficient transformation and gene-editing strategy for *B. napus*. and used it to obtain the first gene-edited winter-type rapeseed. Winter-type Express617 has been used in numerous studies in recent years (Calderwood et al., 2021; Matar et al., 2021; Schilbert et al., 2023) and has an available reference genome (Lee et al., 2020). The distantly related *WUS* gene from sugar beet enabled the regeneration of transgenic winter-type rapeseed by directly inducing shoots, and avoided the laborious construction of inducible systems for the temporally limited expression of *AtWUS*.

## Experimental procedures

### Plant material and transformation

*Brassica napus* winter inbred line Express617 and spring-type rapeseed Westar were used for all transformation experiments. Hypocotyl transformation was conducted based on De Block et al. (1989) with modifications. Seeds were sterilized for 5 min in 70% EtOH, followed by 20 min in 6% NaOCl, and washed with sterile water. Subsequently, seeds were germinated on germination medium (4.9 g L^-1^ MS salts + vitamins + MES, 30 g L^-1^ sucrose, 7 g L^-1^ phyto agar, pH 5.7) and grown in low light for one week. *Agrobacterium tumefaciens* strain GV3101::pMP90 harboring the binary plasmids were grown overnight in LB medium with the appropriate antibiotics (rifampicin, gentamycin, spectinomycin) at 28°C. Bacteria were pelleted and resuspended in wash buffer supplemented with 10 µM acetosyringone (2.45 g L^-1^ MS salts + vitamins, 500 mg L^-1^ MES, 0.01% (v/v) Silwet L-77, pH 5.7) and the OD_600_ was adjusted to 0.1. For co-transformation, *Agrobacterium* culture harboring the 35S::*BvWUS* vector was mixed in a ratio of 1:1 with *Agrobacterium* culture containing the vector with the gene of interest, whereas for the co-transformations with the binary vector 35S::*AtWUS*, a ratio of 1:9 was used. Hypocotyls were cut into about 1-cm segments, submerged in *Agrobacterium* solution, and incubated for 15 min. Before placing the hypocotyl segments on co-cultivation medium (modified after Cardoza and Stewart, 2003: 4.9 g L^-1^ MS salts + vitamins + MES, 30 g L^-1^ sucrose, 3 g L^-1^ Gelrite, 1 mg L^-1^ 2,4-D, 10 µM acetosyringone), they were dabbed into an empty Petri dish to remove excess *Agrobacterium* solution. Hypocotyls were co-cultivated with *Agrobacterium* for 48 h in the dark. Afterwards, hypocotyls were washed in wash buffer supplemented with 400 mg L^-1^ Timentin. Hypocotyls were then placed on regeneration medium (4.9 g L^-1^ MS salts + vitamins + MES, 20 g L^-1^ sucrose, 7 g L^-1^ phyto agar, 2 mg L^-1^ BAP, 0.01 mg L^-1^ Picloram, 0.01 mg L^-1^ NAA, 5 mg L^-1^ AgNO_3_, 400 mg L^-1^ Timentin, 100 mg L^-1^ kanamycin, pH 5.7) and transferred to fresh medium every 3 weeks. Westar hypocotyls were transferred to regeneration medium with reduced BAP concentration (0.5 mg L^-1^ BAP) after 2 weeks. Hypocotyls were cultured in a tissue-culture room with 16 h light at 20–24°C until shoots appeared. Shoots were then excised and transferred to rooting medium (4.9 g L^-1^ MS salts + vitamins + MES, 20 g L^-1^ sucrose, 3 g L^-1^ Gelrite, 0.2 mg L^-1^ NAA, 0.2 mg L^-1^ IBA, 5 mg L^-1^ AgNO_3_, 400 mg L^-1^ Timentin, 100 mg L^-1^ kanamycin, pH 5.7). When the roots were grown sufficiently, rooted plants were transferred to soil and grown in a climate chamber (16 h light, 20°C). Express617 plants were vernalized at 4°C for 8 weeks (16 h light).

### Vector construction

Information for all mentioned genes can be found in Table S10.

### Overexpression vectors

To construct the overexpression vectors, floral buds from Arabidopsis Col-0 and *B. vulgaris* were isolated and RNA was extracted with the Plant RNA Kit, peqGOLD (VWR). The RNA was treated with DNase I and reverse-transcribed into cDNA with the RevertAid First Strand cDNA Synthesis Kit (ThermoFischer Scientific). Primers with *Bam*HI and *Eco*RI restriction sites were used to amplify the ORF of the *AtWUS* and *BvWUS* genes by PCR with Phusion^™^ High-Fidelity DNA Polymerase.

Fragments were blunt-end cloned into pJET1.2 (CloneJET PCR Cloning Kit, ThermoFischer Scientific), and sequences were validated by Sanger sequencing (IKMB, Institute for Clinical Molecular Biology, Kiel, Germany). pJET1.2 containing the correct sequences and overexpression vector B425-p9o-LH-35s-ocs (DNA Cloning Service, Hamburg) were then digested with the respective restriction enzymes and ORFs were ligated into the overexpression vector. Vectors were verified by restriction digestion, to generate the vectors B425-*AtWUS* and B425-*BvWUS*. Both vectors were transformed into *A. tumefaciens* strain GV3101::pMP90.

### zCas9i cloning

The zCas9i cloning kit (Stuttmann et al., 2021) was used to assemble CRISPR vectors. Target sites were designed with the help of CRISPR-P 2.0 (Liu et al., 2017) and checked against the reference genomes of Express617 (Lee et al., 2020) and Westar (Song et al., 2020). Target sites and oligonucleotides are listed in Table S1 and were ordered from Eurofins Genomics. Oligonucleotides were annealed and ligated into shuttle vectors and sgRNAs were assembled into pDGE652 as described by Stuttmann et al. (2021). Plasmids were validated by restriction digestion and Sanger sequencing.

### GUS histochemical assay

Co-transformation was carried out with GV3101::pMP90 harboring the binary plasmid pBin19::pSH4 (Holtorf et al., 1995) with the *uidA* gene under the control of the CaMV 35S promoter. For the GUS assay, hypocotyls or leaf samples were submerged into GUS-substrate solution (1× PBS buffer, 30 µL X-Gluc solution (0.03 g X-Gluc in 300 µL DMSO), 2 µL (v/v) Triton X100) and incubated at 37°C overnight. The tissue was destained with 70% ethanol.

### Genotyping

DNA was isolated after a modified protocol from Dellaporta et al. (1983). To confirm the presence of the integrated transgenes, PCR was conducted with primers binding within the *BvWUS* transgene or in the *nptII* gene of the CRISPR vectors. CRISPR-positive plants were further checked for editing in the respective genes. All PCRs for amplicon sequencing were carried out with Q5 polymerase (NEB).

Therefore, gene-specific primers spanning the target sites were used and with these amplicons, nested PCR was conducted with primers carrying barcodes for pooling of samples. Up to five plants per sample were pooled and amplicons were sequenced by amplicon EZ (Genewiz). For the analysis of pooled sequences, paired-end reads were merged by fastp (Chen, 2023) with standard settings for paired-end reads and the --dont_eval_duplication parameter. Reads were then demultiplexed by using a grep command strategy in Unix. Demultiplexed reads were used for CRISPResso2 analysis (Clement et al., 2019). CRISPRessoPooled was run in mixed mode with previously merged single-end reads with the following parameters and settings: --min_reads_to_use_region 5 --demultiplex_only_at_amplicons --bam_output --min_frequency_alleles_around_cut_to_plot 2 --allele_plot_pcts_only_for_assigned_reference --default_min_aln_score 0 –flexiguide_seq –flexiguide_homology 80. Alignments were manually checked in IGV (Robinson et al., 2011).

## Author Contributions

K.I. carried out the experiments. K.I. and S.M. contributed to experimental design, data analysis, and manuscript writing.

## Acknowledgements

We gratefully acknowledge Bettina Braun for perfect technical assistance. We thank John Chandler for critical reading of the manuscript and helpful suggestions. We thank the Institute of Clinical Molecular Biology in Kiel for providing Sanger sequencing as supported in part by the DFG Clusters of Excellence “Precision Medicine in Chronic Inflammation” and “ROOTS”. This work was supported by the Bundesministerium für Bildung und Forschung (BMBF) grants Enable (FKZ: 031B0801D) and Epibrass (FKZ:031B1223B) to SM.

## Short legends for Supporting Information

Supplementary Figure 1. Evolutionary relationships of taxa.

Supplementary Figure 2. Express617 *bna.clv3* T_1_ seedlings exhibiting red fluorescence from the FAST marker.

Supplementary Figure 3. CRISPResso2 results for *Bna.SPL9.C04b* in T_0_ plants.

Supplementary Figure 4. CRISPResso2 results for *Bna.SPL9.C04a* in T_0_ plants.

Supplementary Figure 5. Overview and timeline of the *BvWUS* co-transformation procedure.

Supplementary Table 1: Target sites for *Bna.CLV3* and *Bna.SPL9/15*

Supplementary Table 2: Gene editing in T_0_ Westar and Express617 *bna.clv3* mutants

Supplementary Table 3: T_1_ progeny of plant *clv3_W2*

Supplementary Table 4: T_1_ progeny of plant *clv3_W7*

Supplementary Table 5: T_1_ progeny of plant *clv3_W8*

Supplementary Table 6: T_1_ progeny of plant *clv3_W10*

Supplementary Table 7: Gene editing in two T_0_ Westar *bna.spl9/15* mutants

Supplementary Table 8: Gene editing in T_1_ Westar progeny of *bna.spl9/15*_W2 and *bna.spl9/15*_W6

Supplementary Table 9: Primers used in this study.

## Notes

This work is original and has not been published elsewhere, nor is it currently under consideration for publication elsewhere. We have also no conflicts of interest to disclose.

### Competing Interest Statement

The authors have declared no competing interest.

