## Supplemental Figures for "Efficient and Versatile Rapeseed Transformation for New Breeding Technologies"

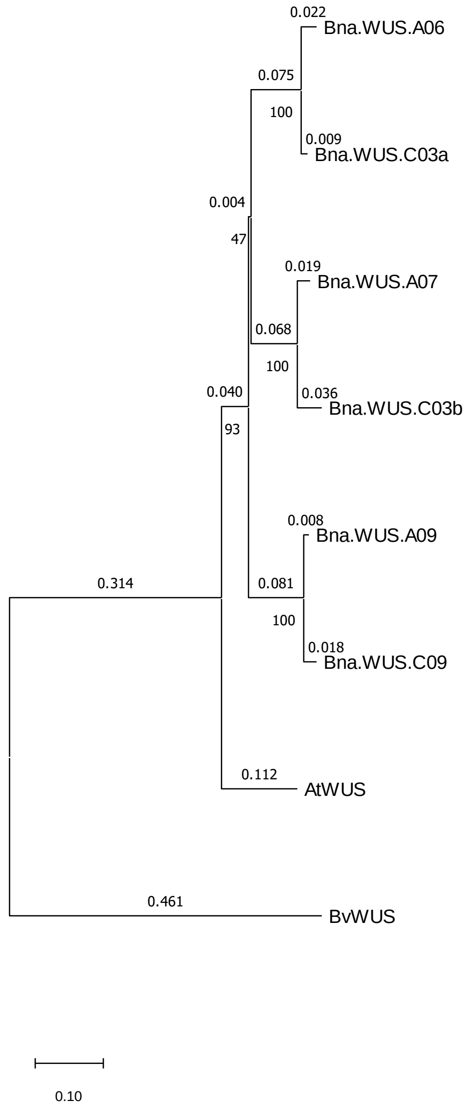


Supplementary Figure 1. Evolutionary relationships of taxa.

The evolutionary history was inferred using the Neighbor-Joining method (Saitou and Nei, 1987). The optimal tree is shown. The percentage of replicate trees in which the associated taxa clustered together in the bootstrap test (500 replicates) is shown below the branches (Felsenstein, 1985). The tree is drawn to scale, with branch lengths (above the branches) in the same units as those of the evolutionary distances used to infer the phylogenetic tree. The evolutionary distances were computed using the Poisson correction method (Zuckerkandl and Pauling, 1965) and are in the units of the number of amino-acid substitutions per site. This analysis involved eight amino-acid sequences. All ambiguous positions were removed for each sequence pair (pairwise deletion option). The final dataset contained 351 positions. Evolutionary analyses were conducted in MEGA11 (Tamura et al., 2021).

Felsenstein, J. (1985). Confidence Limits on Phylogenies: An Approach Using the Bootstrap. *Evolution* 39, 783–791. doi: 10.2307/2408678

Saitou, N., and Nei, M. (1987). The neighbor-joining method: a new method for reconstructing phylogenetic trees. *Molecular Biology and Evolution* 4, 406–425. doi: 10.1093/oxfordjournals.molbev.a040454

Tamura, K., Stecher, G., and Kumar, S. (2021). MEGA11: Molecular Evolutionary Genetics Analysis Version 11. *Molecular Biology and Evolution* 38, 3022–3027. doi: 10.1093/molbev/msab120

Zuckerkandl, E., and Pauling, L. (1965). “Evolutionary Divergence and Convergence in Proteins,” in *Evolving Genes and Proteins*, eds. V. Bryson and H. J. Vogel (Academic Press), 97–166. doi: 10.1016/B978-1-4832-2734-4.50017-6

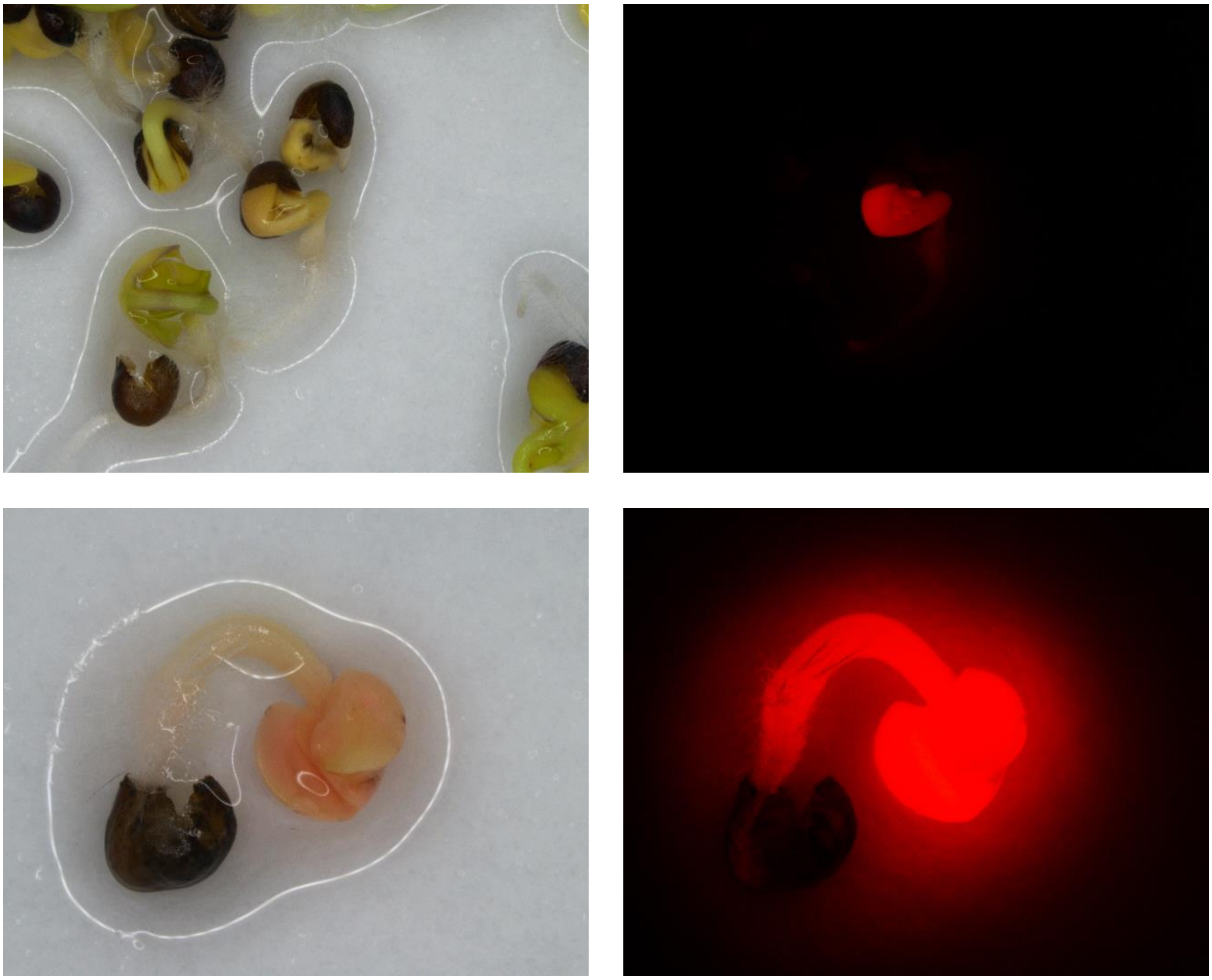

Supplementary Figure 2. Express617 *bna.clv3* $T_1$ seedlings exhibiting red fluorescence from the FAST marker.

*Bna.SPL9.C04b*

*spl9/15*_W1

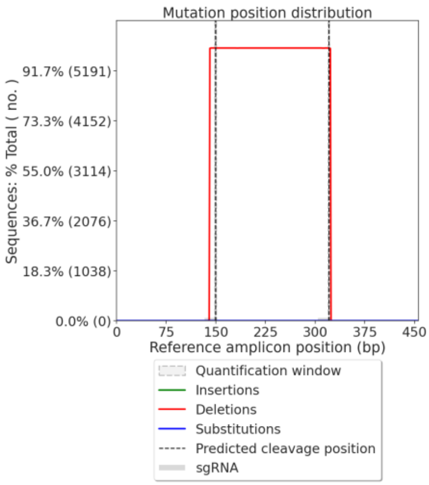


*spl9/15*_W2

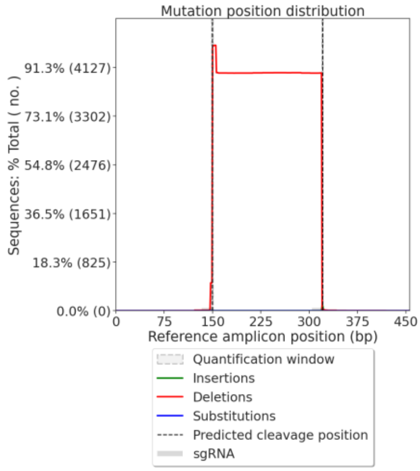


*spl9/15*_W4

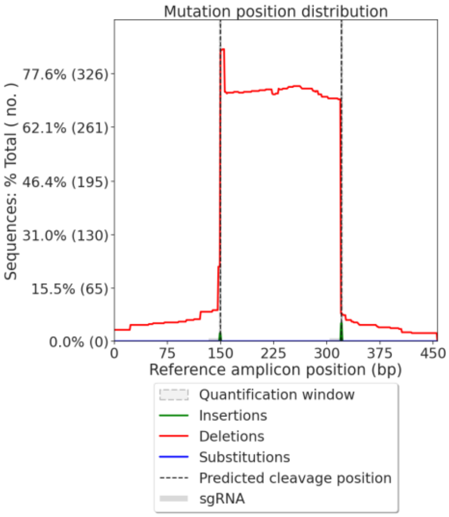


*spl9/15*_W6

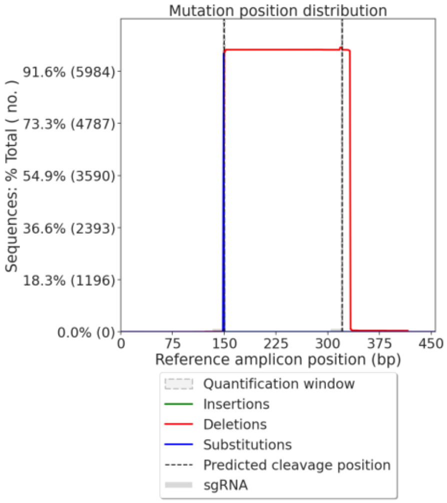


Supplementary Figure 3. CRISPResso2 results for *Bna.SPL9.C04b* in $T_0$ plants *spl9/15*_W1, *spl9/15*_W2, *spl9/15*_W4, and *spl9/15*_W6. Frequency of insertions, deletions, and substitutions across the entire amplicon, considering only modifications that overlap with the quantification window. In all four analyzed plants, deletions between the two target sites occurred.

*Bna.SPL9.C04a*

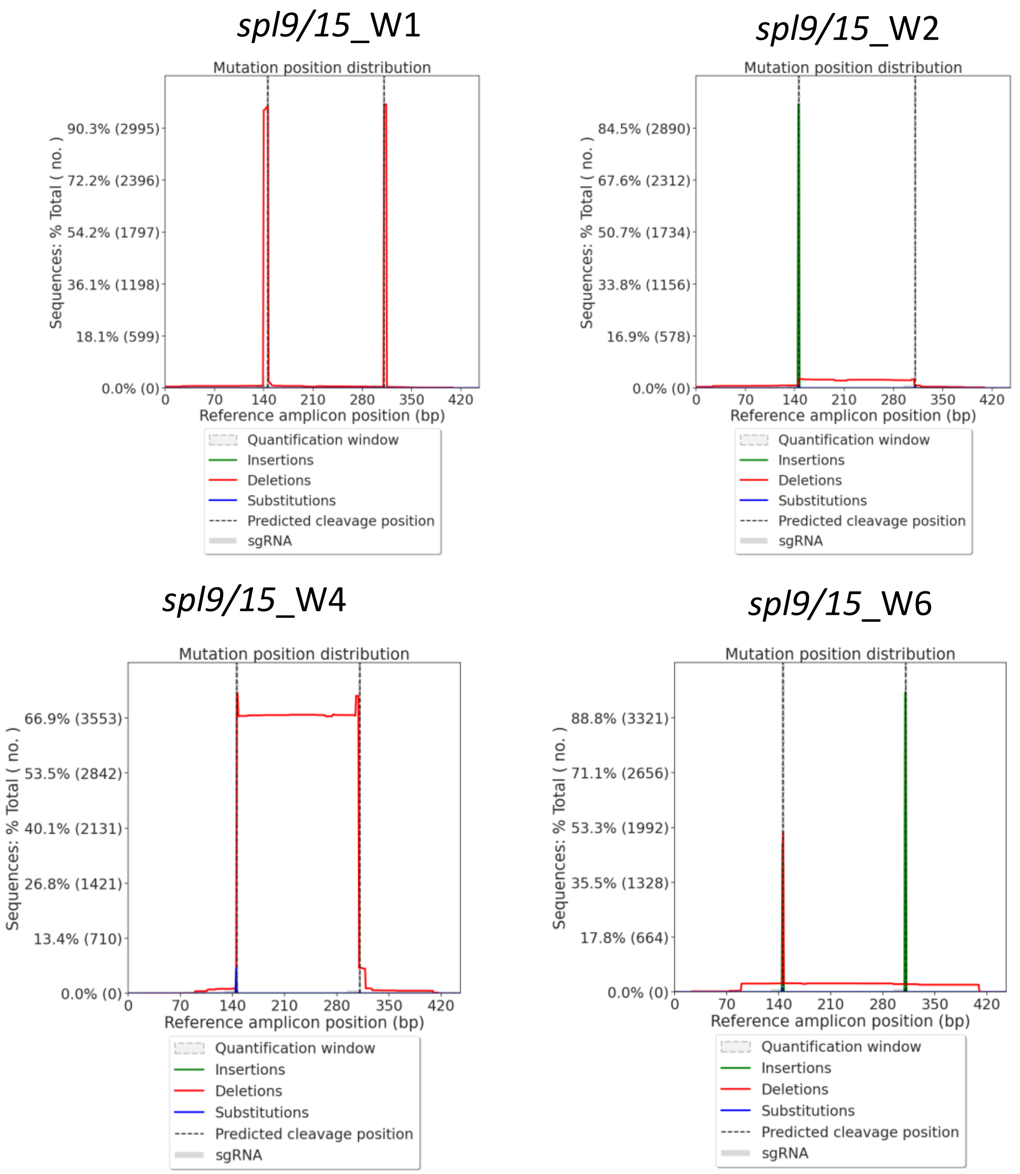


Supplementary Figure 4. CRISPResso2 results for *Bna.SPL9.C04a* in $T_0$ plants *spl9/15*_W1, *spl9/15*_W2, *spl9/15*_W4, and *spl9/15*_W6. Frequency of insertions, deletions, and substitutions across the entire amplicon, considering only modifications that overlap with the quantification window. In all four analyzed plants, the target site with mismatches was edited.

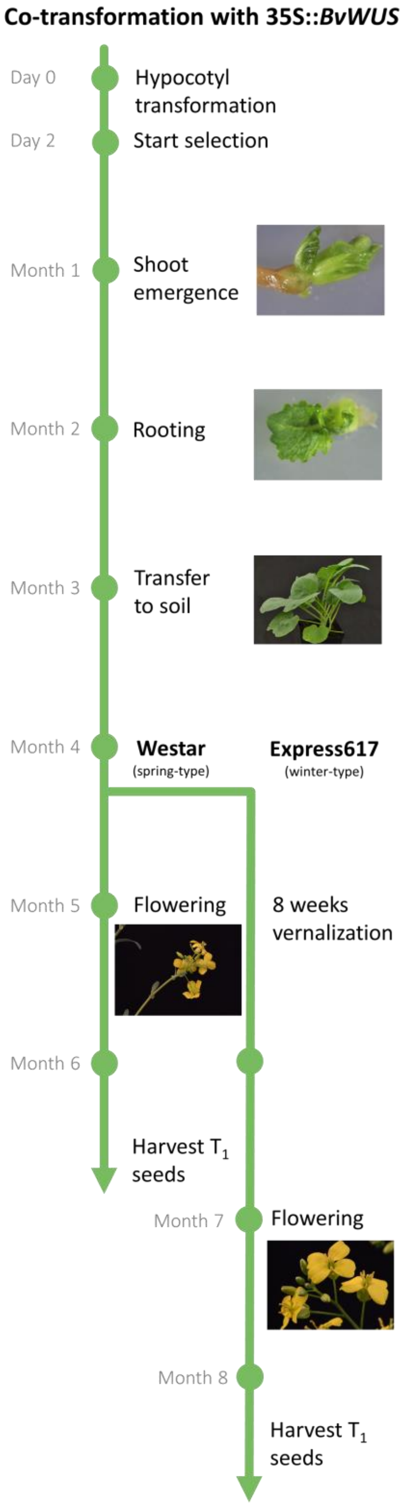


Supplementary Figure 5. Overview and timeline of the *BvWUS* co-transformation procedure.
