## Supplemental Tables for "Efficient and Versatile Rapeseed Transformation for New Breeding Technologies"

Supplementary Table 1: Target sites for *Bna.CLV3* and *Bna.SPL9/15*

| Target gene | sgRNA | Target sequence | PAM | Oligos for sgRNA construction |
|---|---|---|---|---|
| *Bna.CLV3.A04* | TS1 | CTCTCTACAAAATGGATTCG | AGG | attgCTCTCTACAAAATGGATTCG |
| *Bna.CLV3.C04* | TS1 | CTCTCTACAAAATGGATTCG | AGG | aaacCGAATCCATTTTGTAGAGAG |
| *Bna.CLV3.C04* | TS2 | ATGTTCCTGCATGGTGCTTC | TGG | attgATGTTCCTGCATGGTGCTTC |
| | | | | aaacGAAGCACCATGCAGGAACAT |
| *Bna.CLV3.A04* | TS3 | ATGTTCCTGCATGATGCTTC | TGG | attgATGTTCCTGCATGATGCTTC |
| | | | | aaacGAAGCATCATGCAGGAACAT |
| *Bna.CLV3.A04* | TS4 | TCATGCACTTCCCATTCGCA | AGG | attgTCATGCACTTCCCATTCGCA |
| *Bna.CLV3.C04* | TS4 | TCATGCACTTCCCATTCGCA | AGG | aaacTGCGAATGGGAAGTGCATGA |
| *Bna.SPL15.A07* | TS1 | CAGAACCAGCCCGAGTCGGC | TGG | attgCAGAACCAGCCCGAGTCGGC |
| *Bna.SPL15.C06* | TS1 | CAGAACCAGCCCGAGTCGGC | TGG | aaacGCCGACTCGGGCTGGTTCTG |
| *Bna.SPL9.A05* | TS2 | CCAGGCCAGACAGAGTCGGG | TGG | attgCCAGGCCAGACAGAGTCGGG |
| *Bna.SPL9.C04b* | TS2 | CCAGGCCAGACAGAGTCGGG | TGG | aaacCCCGACTCTGTCTGGCCTGG |
| *Bna.SPL9.A04* | TS3 | CCGGGTCAGGCAGAGTCCGG | TGG | attgCCGGGTCAGGCAGAGTCCGG |
| *Bna.SPL9.C04a* | TS3 | CCTGGTCAGGCAGAGTCCGG | TGG | aaacCCGGACTCTGCCTGACCCGG |
| *Bna.SPL15.A07* | TS4 | ACAGCTAGGTGCCAAGTGGA | AGG | attgACAGCTAGGTGCCAAGTGGA |
| *Bna.SPL15.C06* | TS4 | ACAGCTAGGTGCCAAGTGGA | AGG | aaacTCCACTTGGCACCTAGCTGT |
| *Bna.SPL15.A04* | TS4 | GGAGCTAGGTGCCAAGTGGA | AGG | |
| *Bna.SPL15.C04* | TS4 | GCAGCTAGGTGCCAAGTGGA | AGG | |
| *Bna.SPL9.C04b* | TS5 | ATACCAAGATGTCAAGTGGA | AGG | attgATACCAAGATGTCAAGTGGA |
| *Bna.SPL9.A05* | TS5 | ATACCAAGGTGTCAAGTGGA | AGG | aaacTCCACTTGACATCTTGGTAT |
| *Bna.SPL9.A04* | TS6 | ATACCAAGGTGCCAAGTGGA | AGG | attgATACCAAGGTGCCAAGTGGA |
| *Bna.SPL9.C04a* | TS6 | ATACCAAGGTGCCAAGTGGA | AGG | aaacTCCACTTGGCACCTTGGTAT |

Supplementary Table 2: Gene editing in $T_0$ Westar and Express617 *bna.clv3* mutants

| **$T_0$ Plant** | **Allele** | **TS1** | **TS2/3** | **TS4** |
|---|---|---|---|---|
| ***clv3*_W2** | A04.1 | +T | WT | WT |
| | A04.2 | -185 bp | | |
| | C04.1 | -5 bp | WT | +T |
| | C04.2 | +T | WT | WT |
| ***clv3*_W7** | A04.1 | +A | - 133 bp | |
| | A04.2 | - 54 bp | | WT |
| | C04.1 | +G | - 125 bp | |
| | C04.2 | - 11 bp | +A | +A |
| ***clv3*_W8** | A04.1 | - 11 bp | WT | WT |
| | A04.2 | +T | WT | WT |
| | C04.1 | -7 bp | +T | WT |
| | C04.2 | inv | WT | WT |
| ***clv3*_W10** | A04.1 | WT | WT | WT |
| | A04.2 | WT | WT | +A |
| | C04.1 | WT | WT | WT |
| | C04.2 | +A | WT | +G |
| ***clv3*_E2** | A04.1 | - 185 bp | | |
| | A04.2 | - 185 bp | | |
| | C04.1 | - 53 bp | | +G |
| | C04.2 | - 53 bp | | +G |
| ***clv3*_E4** | A04.1 | - 185 bp | | |
| | A04.2 | - 185 bp | | |
| | C04.1 | - 53 bp | | +C |
| | C04.2 | - 53 bp | | +C |
| ***clv3*_E7** | A04.1 | +T | +T | +A |
| | A04.2 | WT | - 125 bp | |
| | C04.1 | WT | WT | WT |
| | C04.2 | +A | WT | +T |
| ***clv3*_E8** | A04.1 | +A | WT | -1 bp |
| | A04.2 | -12 bp, subs. | +A | +A |
| | C04.1 | -43 bp | | +A |
| | C04.2 | +C | +A | WT |
| ***clv3*_E9** | A04.1 | +A | WT | WT |
| | A04.2 | WT | +A | WT |
| | C04.1 | WT | +A | WT |
| | C04.2 | +A | +T | WT |
| ***clv3*_E12** | A04.1 | WT | WT | WT |
| | A04.2 | WT | -4 bp | WT |
| | C04.1 | WT | +A | WT |
| | C04.2 | WT | WT | WT |

### Supplementary Table 3: $T_1$ progeny of plant *clv3*_W2

| $T_1$ Plant | *WUS* | CRISPR |
|---|---|---|
| *clv3*_W2.1 | + | + |
| *clv3*_W2.2 | + | + |
| *clv3*_W2.3 | + | + |
| *clv3*_W2.4 | + | + |
| *clv3*_W2.5 | + | + |
| *clv3*_W2.6 | + | + |
| *clv3*_W2.7 | + | + |
| *clv3*_W2.8 | + | + |
| *clv3*_W2.9 | + | + |
| *clv3*_W2.10 | + | + |
| *clv3*_W2.11 | + | + |
| *clv3*_W2.12 | + | + |
| *clv3*_W2.13 | + | + |
| *clv3*_W2.14 | + | + |
| *clv3*_W2.15 | + | + |
| *clv3*_W2.16 | + | + |
| *clv3*_W2.17 | + | + |
| *clv3*_W2.18 | + | + |
| *clv3*_W2.19 | + | + |
| *clv3*_W2.20 | + | + |
| *clv3*_W2.21 | + | + |
| *clv3*_W2.22 | + | + |
| *clv3*_W2.23 | + | + |
| *clv3*_W2.24 | + | + |
| *clv3*_W2.25 | + | + |
| *clv3*_W2.26 | + | + |
| *clv3*_W2.27 | + | + |
| *clv3*_W2.28 | + | + |
| *clv3*_W2.29 | + | + |
| *clv3*_W2.30 | + | + |
| *clv3*_W2.31 | + | + |
| *clv3*_W2.32 | + | + |
| *clv3*_W2.33 | + | + |
| *clv3*_W2.34 | + | + |
| *clv3*_W2.35 | + | + |
| *clv3*_W2.36 | + | + |
| *clv3*_W2.37 | + | + |
| *clv3*_W2.38 | + | + |
| *clv3*_W2.39 | + | + |
| *clv3*_W2.40 | + | + |
| *clv3*_W2.41 | + | + |
| *clv3*_W2.42 | + | + |
| *clv3*_W2.43 | + | + |
| *clv3*_W2.44 | + | + |
| *clv3*_W2.45 | + | + |
| *clv3*_W2.46 | + | + |
| *clv3*_W2.47 | + | + |
| *clv3*_W2.48 | + | + |

Supplementary Table 4: $T_1$ progeny of plant *clv3*_W7

| $T_1$ Plant | ***WUS*** | **CRISPR** |
|---|---|---|
| *clv3*_W7.1 | + | + |
| *clv3*_W7.2 | - | + |
| *clv3*_W7.3 | + | + |
| *clv3*_W7.4 | + | + |
| *clv3*_W7.5 | + | + |
| *clv3*_W7.6 | + | + |
| *clv3*_W7.7 | + | + |
| *clv3*_W7.8 | + | + |
| *clv3*_W7.9 | + | + |
| *clv3*_W7.10 | - | + |
| *clv3*_W7.11 | + | + |
| *clv3*_W7.12 | + | + |
| *clv3*_W7.13 | + | + |
| *clv3*_W7.14 | + | + |
| *clv3*_W7.15 | - | + |
| *clv3*_W7.16 | + | + |
| *clv3*_W7.17 | + | + |
| *clv3*_W7.18 | + | + |
| *clv3*_W7.19 | + | + |
| *clv3*_W7.20 | + | + |
| *clv3*_W7.21 | + | + |
| *clv3*_W7.22 | + | + |
| *clv3*_W7.23 | + | + |
| *clv3*_W7.24 | + | + |
| *clv3*_W7.25 | + | + |
| *clv3*_W7.26 | - | + |
| *clv3*_W7.27 | - | + |
| *clv3*_W7.28 | + | + |
| *clv3*_W7.29 | - | + |
| *clv3*_W7.30 | + | + |
| *clv3*_W7.31 | + | + |
| *clv3*_W7.32 | + | + |
| *clv3*_W7.33 | + | + |
| *clv3*_W7.34 | - | + |
| *clv3*_W7.35 | + | + |
| *clv3*_W7.36 | + | + |
| *clv3*_W7.37 | + | + |
| *clv3*_W7.38 | + | + |
| *clv3*_W7.39 | - | + |
| *clv3*_W7.40 | + | + |
| *clv3*_W7.41 | - | + |
| *clv3*_W7.42 | + | + |
| *clv3*_W7.43 | + | + |
| *clv3*_W7.44 | + | + |
| *clv3*_W7.45 | + | + |
| *clv3*_W7.46 | + | + |
| *clv3*_W7.47 | + | + |
| *clv3*_W7.48 | + | + |

Supplementary Table 5: $T_1$ progeny of plant *clv3*_W8

| $T_1$ Plant | *WUS* | CRISPR |
|---|---|---|
| *clv3*_W8.1 | - | + |
| *clv3*_W8.2 | - | + |
| *clv3*_W8.3 | - | + |
| *clv3*_W8.4 | - | + |
| *clv3*_W8.5 | - | - |
| *clv3*_W8.6 | - | - |
| *clv3*_W8.7 | - | + |
| *clv3*_W8.8 | - | + |
| *clv3*_W8.9 | - | + |
| *clv3*_W8.10 | - | + |
| *clv3*_W8.11 | - | + |
| *clv3*_W8.12 | - | + |
| *clv3*_W8.13 | - | + |
| *clv3*_W8.14 | - | + |
| *clv3*_W8.15 | - | + |
| *clv3*_W8.16 | - | + |
| *clv3*_W8.17 | - | - |
| *clv3*_W8.18 | - | + |
| *clv3*_W8.19 | - | + |
| *clv3*_W8.20 | - | + |
| *clv3*_W8.21 | - | + |
| *clv3*_W8.22 | - | + |
| *clv3*_W8.23 | - | + |
| *clv3*_W8.24 | - | + |
| *clv3*_W8.25 | - | + |
| *clv3*_W8.26 | - | + |
| *clv3*_W8.27 | - | + |
| *clv3*_W8.28 | - | + |
| *clv3*_W8.29 | - | + |
| *clv3*_W8.30 | - | + |
| *clv3*_W8.31 | - | + |
| *clv3*_W8.32 | - | + |
| *clv3*_W8.33 | - | + |
| *clv3*_W8.34 | - | + |
| *clv3*_W8.35 | - | + |
| *clv3*_W8.36 | - | + |
| *clv3*_W8.37 | - | + |
| *clv3*_W8.38 | - | + |
| *clv3*_W8.39 | - | + |
| *clv3*_W8.40 | - | + |
| *clv3*_W8.41 | - | + |
| *clv3*_W8.42 | - | + |
| *clv3*_W8.43 | - | + |
| *clv3*_W8.44 | - | + |
| *clv3*_W8.45 | - | + |
| *clv3*_W8.46 | - | + |
| *clv3*_W8.47 | - | + |
| *clv3*_W8.48 | - | + |

Supplementary Table 6: $T_1$ progeny of plant *clv3*_W10

| $T_1$ Plant | *WUS* | CRISPR |
|---|---|---|
| *clv3*_W10.1 | - | + |
| *clv3*_W10.2 | - | + |
| *clv3*_W10.3 | - | + |
| *clv3*_W10.4 | - | - |
| *clv3*_W10.5 | - | + |
| *clv3*_W10.6 | - | + |
| *clv3*_W10.7 | - | - |
| *clv3*_W10.8 | - | - |
| *clv3*_W10.9 | - | - |
| *clv3*_W10.10 | - | + |
| *clv3*_W10.11 | - | - |
| *clv3*_W10.12 | - | + |
| *clv3*_W10.13 | - | + |
| *clv3*_W10.14 | - | + |
| *clv3*_W10.15 | - | - |
| *clv3*_W10.16 | - | + |
| *clv3*_W10.17 | - | + |
| *clv3*_W10.18 | - | + |
| *clv3*_W10.19 | - | + |
| *clv3*_W10.20 | - | - |
| *clv3*_W10.21 | - | + |
| *clv3*_W10.22 | - | + |
| *clv3*_W10.23 | - | + |
| *clv3*_W10.24 | - | - |
| *clv3*_W10.25 | - | - |
| *clv3*_W10.26 | - | - |
| *clv3*_W10.27 | - | + |
| *clv3*_W10.28 | - | - |
| *clv3*_W10.29 | - | + |
| *clv3*_W10.30 | - | + |
| *clv3*_W10.31 | - | + |
| *clv3*_W10.32 | - | - |
| *clv3*_W10.33 | - | - |
| *clv3*_W10.34 | - | + |
| *clv3*_W10.35 | - | - |
| *clv3*_W10.36 | - | + |
| *clv3*_W10.37 | - | + |
| *clv3*_W10.38 | - | + |
| *clv3*_W10.39 | - | + |
| *clv3*_W10.40 | - | - |
| *clv3*_W10.41 | - | + |
| *clv3*_W10.42 | - | + |
| *clv3*_W10.43 | - | - |
| *clv3*_W10.44 | - | + |
| *clv3*_W10.45 | - | + |
| *clv3*_W10.46 | - | + |
| *clv3*_W10.47 | - | + |

### Supplementary Table 7: Gene editing in two $T_0$ Westar *bna.spl9/15* mutants

| **$T_0$ Plant** | **Gene** | **Allele** | **TS2** | **TS6** |
|---|---|---|---|---|
| ***spl9/15*_W2** | *SPL9* | A04.1 | T ins | WT |
| | | A04.2 | G ins | WT |
| | | | **TS2** | **TS5** |
| | | A05.1 | A ins | WT |
| | | A05.2 | C ins | WT |
| | | | **TS3** | **TS6** |
| | | C04a.1 | T ins | WT |
| | | C04a.2 | T ins | WT |
| | | | **TS2** | **TS5** |
| | | C04b.1 | 170 bp del | |
| | | C04b.2 | 9 bp del | WT |
| | | | **TS1** | **TS4** |
| | *SPL15* | A07.1 | T ins | 2 bp del |
| | | A07.2 | A ins | T ins |
| | | C06.1 | T ins | T ins |
| | | C06.2 | T ins | T ins |
| | | | | **TS4** |
| | | A04.1 | | T ins |
| | | A04.2 | | 3 bp del |
| | | C04.1 | | A ins |
| | | C04.2 | | T ins |
| | | | **TS2** | **TS6** |
| ***spl9/15*_W6** | *SPL9* | A04.1 | T ins | T ins |
| | | A04.2 | A ins | T ins |
| | | | **TS2** | **TS5** |
| | | A05.1 | G ins | A ins |
| | | A05.2 | T ins | T ins |
| | | | **TS3** | **TS6** |
| | | C04a.1 | C del | T ins |
| | | C04a.2 | A ins | T ins |
| | | | **TS2** | **TS5** |
| | | C04b.1 | T ins + 183 bp del | |
| | | C04b.2 | G del | 3 bp del |
| | | | **TS1** | **TS4** |
| | *SPL15* | A07.1 | 157 bp del | |
| | | A07.2 | WT | T ins |
| | | C06.1 | T ins | T ins |
| | | C06.2 | T ins | T ins |
| | | | | **TS4** |
| | | A04.1 | | WT |
| | | A04.2 | | T ins |
| | | C04.1 | | 4 bp del |
| | | C04.2 | | T ins |

### Supplementary Table 8: Gene editing in $T_1$ Westar progeny of *bna.spl9/15*_W2 and *bna.spl9/15*_W6

| ***spl9/15*_W2.13** | | |
|---|---|---|
| **SPL9.A04** | **TS2** | **TS6** |
| | T ins | WT |
| | G ins | WT |
| **SPL9.A05** | **TS2** | **TS5** |
| | A ins | WT |
| | A ins | WT |
| **SPL9.C04a** | **TS3** | **TS6** |
| | T ins | WT |
| | T ins | WT |
| **SPL9.C04b** | **TS2** | **TS5** |
| | 170 bp del | |
| | 9 bp del | WT |
| **SPL15.A07** | **TS1** | **TS4** |
| | A ins | T ins |
| | A ins | T ins |
| **SPL15.C06** | **TS1** | **TS4** |
| | T ins | T ins |
| | T ins | T ins |
| **SPL15.A04** | | **TS4** |
| | | T ins |
| | | 3 bp del |
| **SPL15.C04** | | **TS4** |
| | | T ins |
| | | T ins |

| ***spl9/15*_W2.26** | | |
|---|---|---|
| **SPL9_A04** | **TS2** | **TS6** |
| | T ins | WT |
| | G ins | WT |
| **SPL9_A05** | **TS2** | **TS5** |
| | C ins | WT |
| | C ins | WT |
| **SPL9_C04a** | **TS3** | **TS6** |
| | T ins | WT |
| | T ins | WT |
| **SPL9_C04b** | **TS2** | **TS5** |
| | 170 bp del | |
| | 9 bp del | WT |
| **SPL15_A07** | **TS1** | **TS4** |
| | T ins | 2 bp del |
| | A ins | T ins |
| **SPL15_C06** | **TS1** | **TS4** |
| | T ins | T ins |
| | T ins | T ins |
| **SPL15_A04** | | **TS4** |
| | | T ins |
| | | T ins |
| **SPL15_C04** | | **TS4** |
| | | T ins |
| | | T ins |

| ***spl9/15*_W6.21** | | |
|---|---|---|
| **SPL9_A04** | **TS2** | **TS6** |
| | T ins | T ins |
| | A ins | T ins |
| **SPL9_A05** | **TS2** | **TS5** |
| | G ins | A ins |
| | T ins | T ins |
| **SPL9_C04a** | **TS3** | **TS6** |
| | C del | T ins |
| | C del | T ins |
| **SPL9_C04b** | **TS2** | **TS5** |
| | T ins + 183 bp del | |
| | T ins + 183 bp del | |
| **SPL15_A07** | **TS1** | **TS4** |
| | 157 bp del | |
| | WT | T ins |
| **SPL15_C06** | **TS1** | **TS4** |
| | T ins | T ins |
| | T ins | T ins |
| **SPL15_A04** | | **TS4** |
| | | T ins |
| | | T ins |
| **SPL15_C04** | | **TS4** |
| | | T ins |
| | | T ins |

| ***spl9/15*_W6.32** | | |
|---|---|---|
| **SPL9_A04** | **TS2** | **TS6** |
| | A ins | T ins |
| | A ins | T ins |
| **SPL9_A05** | **TS2** | **TS5** |
| | G ins | A ins |
| | T ins | T ins |
| **SPL9_C04a** | **TS3** | **TS6** |
| | C del | T ins |
| | A ins | T ins |
| **SPL9_C04b** | **TS2** | **TS5** |
| | T ins + 183 bp del | |
| | T ins + 183 bp del | |
| **SPL15_A07** | **TS1** | **TS4** |
| | 157 bp del | |
| | WT | T ins |
| **SPL15_C06** | **TS1** | **TS4** |
| | T ins | T ins |
| | T ins | T ins |
| **SPL15_A04** | | **TS4** |
| | | T ins |
| | | T ins |
| **SPL15_C04** | | **TS4** |
| | | T ins |
| | | T ins |

### Supplementary Table 9: Primers used in this study.

| Primer | Sequence |
|---|---|
| AtWUS_BamHI_for | AGTTGGATCCATGGAGCCGCCACAGCATCA |
| AtWUS_EcoRI_rev | ATTCGAATTCTAGTTCAGACGTAGCTCAA |
| BvWUS_BamHI_for | AGTTGGATCCATGAGTAATACTACAAGTAG |
| BvWUS_EcoRI_rev | ATTCGAATTCCTAAAGATTTTGTGAATAAC |
| AtWUS_for | AGCCGATCAGATCCAGAAGA |
| AtWUS_rev | AACCGAGTTGGGTGATGAAG |
| BvWUS_for | TTCCTCTCTTCCCAATGCAC |
| BvWUS_rev | AGGGAGGTATTCCCACCATT |
| Kan1-r | CTTCCCGCTTCAGTGACAAC |
| Kan2-f | TTGGGTGGAGAGGCTATTCG |
| CLV3_A04_for | GTGTTCTATATTCCGGACATACG |
| CLV3_A04_rev | CTGAAGGGACAGTCCTTAGT |
| CLV3_C04_for | GGAGAAAGGATCTAGTGATCG |
| CLV3_C04_rev | GCTAAGGACTGTCCCTTCAG |
| NGS_CLV3_for | CTTGCAGCCTATAAATGATTGC |
| NGS_CLV3_rev | AACACGAGATAGATGTCCG |
| SPL9_A04_for | CACAGTTGGTTGATAAGCATTTAG |
| SPL9_A04_rev | CAGACCGTGTTAGCTTCTAGA |
| SPL9_A05_for | CATGAACCAAGCGATGAGTAC |
| SPL9_A05_rev | CCTGGTCCCATAACATTATGC |
| SPL9_C04a_for | GCTGGAACTGCCTTATGTTG |
| SPL9_C04a_rev | CAGTCCGTGTTAACTTCTAAGG |
| SPL9_C04b_for | GCAAGTACTTAAAGGTCGTAACC |
| SPL9_C04b_rev | ACCATTTCCTGGCCCATG |
| SPL15_A07_for | AGATGTTCACTACAGAAAACG |
| SPL15_A07_rev | ACCTAACCATATAGAGATGGAGAG |
| SPL15_C06_for | GTTCATTCACTGCAGAGAACC |
| SPL15_C06_rev | CATATAGAGATGGAGCGATGTTG |
| NGS_SPL9_A04_for | CGTCCTTTCTTTAAACCAAGACAG |
| NGS_SPL9_A05_for | CCTTTCCTTTAAACCGAGACAG |
| NGS_SPL9_C04a_for | GTCCTTTCTTCAAACCAAGACAG |
| NGS_SPL9_C04b_for | CCTTGCCTTTAAACCGAGACAG |
| NGS_SPL9_rev | GAACCTGCTGCACTGTTGAC |
| NGS_SPL15A04_C04_for | TGGAGTTACTAATGGGTTCGG |
| NGS_SPL15A04_C04_rev | CATTGTTGGCAAAACCTTTGA |
| NGS_SPL15A07_C06_for | TCGGCTGGTTCCTCGTCTA |
| NGS_SPL15A07_C06_rev | CATTGCTGGCAAAACCTTTGG |

### Supplementary Table 10: Gene information

| Species | Genotype | Gene | Gene identifier | Reference genome |
|---|---|---|---|---|
| *A.thaliana* | | *AtWUS* | AT2G17950 | |
| *B.vulgaris* | | *BvWUS* | g68555.t1 | RefBeet1.2 (Dohm et al., 2014) |
| *B.napus* | Express617 | *Bna.CLV3* | A04p016980.1_BnaEXP | Express617 (Lee et al., 2020) |
| | | | C04p038910.1_BnaEXP | |
| *B.napus* | Westar | *Bna.CLV3* | BnaA04T0166900WE | Westar v0 (Song et al., 2020) |
| | | | BnaC04T0465000WE | |
| | | *Bna.SPL9* | BnaA04T0257100WE | |
| | | | BnaC04T0572100WE | |
| | | | BnaC04T0031400WE | |
| | | | BnaA05T0019400WE | |
| | | *Bna.SPL15* | BnaA04T0025800WE | |
| | | | BnaC04T0286900WE | |
| | | | BnaC06T0236300WE | |
| | | | BnaA07T0194600WE | |
